# Multi-Omic Profiling Reveals Dynamics of the Phased Progression of Pluripotency

**DOI:** 10.1101/415430

**Authors:** Pengyi Yang, Sean J. Humphrey, Senthilkumar Cinghu, Rajneesh Pathania, Andrew J. Oldfield, Dhirendra Kumar, Dinuka Perera, Jean Y.H. Yang, David E. James, Matthias Mann, Raja Jothi

## Abstract

Pluripotency is highly dynamic and progresses through a continuum of pluripotent stem-cell states. The two states that bookend the pluripotency continuum, naïve and primed, are well characterized, but our understanding of the intermediate states and transitions between them remain incomplete. Here, we dissect the dynamics of pluripotent state transitions underlying pre-to post-implantation epiblast differentiation. Through comprehensive mapping of the proteome, phosphoproteome, transcriptome, and epigenome of mouse embryonic stem cells transitioning from naïve to primed pluripotency, we find that rapid, acute, and widespread changes to the phosphoproteome precede ordered changes to the epigenome, transcriptome, and proteome. Reconstruction of kinase-substrate networks reveals signaling cascades, dynamics, and crosstalk. Distinct waves of global proteomic changes demarcate discrete phases of pluripotency, characterized by cell-state-specific surface marker expression. Our data provide new insights into the multi-layered control of the phased progression of pluripotency and a foundation for modeling mechanisms underlying pre-to post-implantation epiblast differentiation.

**HIGHLIGHTS:**

- Multi-ome maps of cells transitioning from naïve to primed pluripotency
- Phosphoproteome dynamics precede changes to epigenome, transcriptome, and proteome
- Kinase-substrate network reconstruction uncovers signaling dynamics and crosstalk
- Proteins and cell surface markers that track pluripotent state transitions
- Comparative analysis of mouse and human pluripotent states

## INTRODUCTION

Pluripotency describes the developmental potential of a cell to give rise to derivatives of all three primary germ layers. Although pluripotency is ephemeral *in vivo*, pluripotent stem cells (PSCs), derived from various stages of early embryonic development, can self-renew indefinitely *in vitro* under defined culture conditions while retaining their pluripotent status (Nichols and Smith, 2009). Studies of the early mouse embryo and PSCs in culture have led to the proposition that embryonic pluripotency is highly dynamic and proceeds through a continuum of pluripotent stem-cell states (De Los Angeles et al., 2015; Hackett and Surani, 2014; Nichols and Smith, 2009; Rossant and Tam, 2017; Shahbazi et al., 2017; Weinberger et al., 2016; Wu and Izpisua Belmonte, 2015). At one end of this continuum is the naïve pluripotent state (Nichols and Smith, 2009), sometimes also referred to as the ground state (Hackett and Surani, 2014; Marks et al., 2012; Ying et al., 2008), representing the broadest and most unrestricted developmental potential that exists in the pre-implantation mouse embryo from approximately embryonic day 3.75 (E3.75) to E4.75 (Boroviak et al., 2014). At the other end of this continuum is the primed pluripotent state, representing the developmentally restricted potential that exists in pluripotent cells from post-implantation mouse epiblasts (E5.5-E8.25), which are lineage-primed for differentiation.

Embryonic stem cells (ESCs), derived from the inner cell mass (ICM) of pre-implantation mouse blastocysts (Figure 1A) and maintained under defined culture conditions known as 2i+LIF (Ying et al., 2008), most closely resemble naïve epiblasts of the pre-implantation embryo (Boroviak et al., 2014; Boroviak et al., 2015). Hence, ESCs are considered to capture the naïve pluripotent state. Epiblast stem cells (EpiSCs), isolated from pre-gastrulation (E5.5) to late-bud (E8.25) stage post-implantation mouse epiblasts (Brons et al., 2007; Tesar et al., 2007) (Figure 1A), are developmentally comparable to the late-gastrulation stage (E7.0) embryo, irrespective of the original developmental stage (E5.5-E8.25) of their source tissue (Kojima et al., 2014); these cells are considered archetypal representative of the primed pluripotent state. Interestingly, conventional human ESCs (hESCs), derived from pre-implantation human blastocysts, exhibit molecular and morphological characteristics that are more similar to primed EpiSCs than to naïve ESCs (Davidson et al., 2015; De Los Angeles et al., 2015; Hackett and Surani, 2014; Rossant and Tam, 2017; Weinberger et al., 2016; Wu and Izpisua Belmonte, 2015). Several protocols that reprogram hESCs back to the ground state have been proposed (Chan et al., 2013; Gafni et al., 2013; Takashima et al., 2014; Theunissen et al., 2014; Ware et al., 2014), but they each generate “naïve” hESCs with distinct transcriptional profiles (Davidson et al., 2015; Huang et al., 2014) and fail to recover the naïve epiblast methylation landscape (Pastor et al., 2016). Although these purported naïve hESCs satisfy some features of mouse criteria for the naïve pluripotent state, whether they can be considered equivalent to naïve mouse ESCs remains an open question.

**Figure 1.**
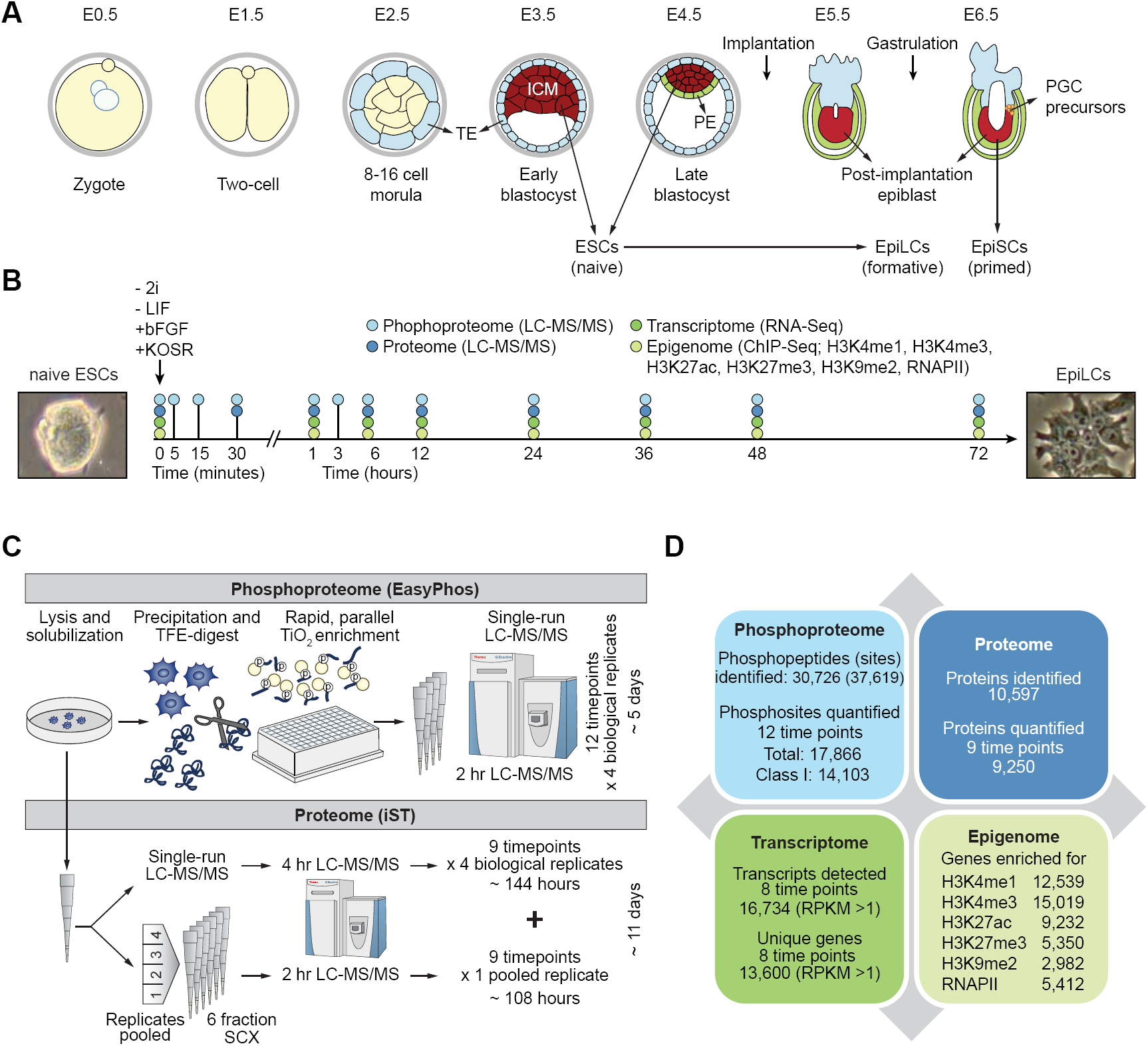
High temporal-resolution profiling of the proteome, phosphoproteome, transcriptome, and epigenome during ESC to EpiLC transition. (A) Developmental events during embryogenesis in mouse embryos. ICM, inner cell mass; ESCs, embryonic stem cells; TE, tropectoderm; PE, primitive endoderm; EpiLC, epiblast-like cells; EpiSCs, epiblast stem cells; PGCs, primordial stem cells. (B) Schematic showing EpiLC induction from ESCs grown in 2i+LIF medium. Proteome, phosphoproteome, transcriptome, and epigenome were profiled at indicated time-points. Phase contrast images correspond to representative ESC colony grown in 2i+LIF medium (left) and cells undergoing morphological changes at 72h post EpiLC induction (right). (C) Schematic of mass spectrometry (MS)-based experimental protocols used for proteome and phosphoproteome profiling. (D) Summary statistics of proteins, phosphosites, transcripts, and epigenetic marks profiled. See also Figure S1 and S2.

ESCs are highly competent to form high-contribution mouse chimeras with germline transmission, following microinjection into pre-implantation embryos. EpiSCs, however, do not integrate well into host blastocysts, likely because they correspond to a developmentally advanced stage compared to the host pre-implantation environment, and thus contribute poorly or not at all to blastocyst chimeras (Dejosez and Zwaka, 2012; Hackett and Surani, 2014; Han et al., 2010; Nichols and Smith, 2009; Weinberger et al., 2016; Wu and Izpisua Belmonte, 2015). Conversely, when grafted into post-implantation (E7.5) embryos in whole embryo culture, EpiSCs but not ESCs efficiently incorporate into the host and contribute to all three germ layers (Huang et al., 2012). Consequently, primed EpiSCs are considered to be functionally and developmentally distinct from naïve epiblast and ESCs (De Los Angeles et al., 2015; Weinberger et al., 2016). While the naïve and primed states, which bookend the pluripotency continuum, are well characterized (Kojima et al., 2014; Marks et al., 2012), our understanding of the intermediate pluripotent states and the transitions between them remain incomplete.

Cell signaling underlies transcriptional and/or epigenetic control of a vast majority of cell fate decisions during early embryonic development (Dejosez and Zwaka, 2012; Hackett and Surani, 2014; Rossant and Tam, 2017; Weinberger et al., 2016). Yet, our understanding of the signaling dynamics during pluripotent state transitions and how they instruct epigenetic and/or transcriptional programs controlling ICM to post-implantation epiblast differentiation remains poorly understood. Recent advances in mass spectrometry (MS)-based proteomics now allow for near-comprehensive characterization of proteomes and phosphorylation events (Aebersold and Mann, 2016; Altelaar et al., 2013; Humphrey et al., 2015; Riley and Coon, 2016; Robles et al., 2017; Sharma et al., 2014). To elucidate the signaling and molecular dynamics that underlie pluripotent state transitions, here we generated comprehensive high-temporal-resolution maps of the phosphoproteome, proteome, transcriptome, and epigenome of embryonic stem cells transitioning from naïve to primed pluripotency. Our data provide new insights into the multi-layered control of the phased progression of pluripotency and a foundation for investigating mechanisms underlying ICM to post-implantation epiblast differentiation.

## RESULTS

### High temporal-resolution maps of the proteome, phosphoproteome, transcriptome, and epigenome of cells transitioning from naïve to primed pluripotency

To elucidate the temporal dynamics of the phosphoproteome, proteome, epigenome, and transcriptome during the transition from naïve to primed pluripotency, we employed a previously validated system to induce naïve mouse ESCs to post-implantation pre-gastrulating epiblast-like cells (EpiLCs) (Buecker et al., 2014; Hayashi et al., 2011; Kurimoto et al., 2015; Shirane et al., 2016), which closely resemble the early post-implantation epiblast (E5.5-E6.5) than do EpiSCs (Hayashi et al., 2011). EpiLCs were induced by plating naïve ESCs, grown in ground state under serum-free 2i+LIF medium, onto fibronectin-coated plates in N2B27 medium containing activin A, bFGF, and knockout serum replacement (KOSR, 1%) (Hayashi et al., 2011). Consistent with previous reports, within 48 h of EpiLC induction, morphological transformation in the form of flattened epithelial structures resembling epiblasts was evident (Figure S1A). RNA analysis using quantitative RT-PCR confirmed downregulation of naïve pluripotency/ICM-associated genes (*Nanog, Klf4, Prdm14*) accompanied by induction of post-implantation epiblast-associated genes (*Fgf5, Otx2, Pou3f1*/*Oct6*) (Figure S1B) (Buecker et al., 2014; Hayashi et al., 2011; Kalkan et al., 2017). Although no dramatic changes in transcript levels of these marker genes were evident after 48 h post induction, we included the 72-h time-point in our analyses to capture changes to the proteome that may lag changes to the transcriptome.

Using advances in MS-based proteomics (Kulak et al., 2014) and our EasyPhos workflow (Humphrey et al., 2015), together with next-generation sequencing, we generated maps of the phosphoproteome, proteome, transcriptome, and epigenome of cells at various time-points during the 72-h ESC to EpiLC transition (Figure 1B). To capture the earliest signaling responses, we profiled the phosphoproteome of transitioning cells at high temporal-resolution within the first hour post-induction (Figure 1B). All MS experiments were performed in biological quadruplicates. In addition, to enhance coverage of the proteome measurements, we pooled the four biological replicates from each time-point and performed StageTip-based Strong Cation Exchange (SCX) fractionation (Wisniewski et al., 2009) of this pooled sample for the proteome runs (Figures 1C, S2A, **and** S2B). All MS data were analyzed using the MaxQuant computational platform (Cox and Mann, 2008; Tyanova et al., 2016).

Our single-run EasyPhos workflow (Humphrey et al., 2015) produced excellent phosphopeptide coverage, quantifying over 15,000 phosphopeptides in every run (Figure S2C). This yielded a total of 30,726 distinct phosphopeptides from which we identified 37,619 individual phosphorylation sites (Figure 1d). Phosphosite localization confidence was high, with >80% (26,180) of the quantified phosphosites accurately localized to a single amino acid (mean localization probability for quantified sites: 0.96) (Figure S2D **and Methods**). A total of 17,866 phosphosites and over half of the Class 1 phosphosites (14,103) were quantified across all 12 time-points analyzed (Figure 1D; **Table S1**). From our proteome runs, we identified over 160,000 distinct peptides and quantified a grand total of 10,597 proteins across all samples and 9,250 proteins in every sample (Figure 1D). Quantification coverage at the proteome level was also very high, with 9,250 proteins quantified across all profiled time-points (Figure 1D; **Table S2**).

Using paired-end RNA-Seq, we mapped the transcriptome across eight time-points during the 72-h time-course and detected a total of 16,734 transcripts (RPKM > 1) corresponding to 13,600 unique genes (Figures 1D **and** S2E**; Table S3**). ChIP-Seq analyses of the chromatin, collected from the same eight time-points, using antibodies against common histone modifications (H3K4me1, H3K4me3, and H3K27ac: associated with the promoters of transcriptionally active genes; H3K27me3 and H3K9me2: associated with the promoters of silent genes) and RNA Polymerase II (RNAPII) identified several thousand transcriptionally active/poised genes (Figures 1D **and** S2F**; Table S4**).

### ESCs exit the naïve pluripotent state by about 36 hours post-induction

Principal component analysis (PCA) and unsupervised hierarchical clustering of the transcriptome, proteome, phosphoproteome, or epigenome revealed clear time-dependent separation of the data (Figures 2A-C **and** S3), with global changes to the phosphoproteome evident as early as five minutes post-induction (Figure 2C), suggesting that the clustering is driven largely by differences in the underlying biological signal across various time-points. PCA analysis of our transcriptomic data, in conjunction with the recently published RNA-Seq data obtained from ESCs transitioning out of naive ground state pluripotency (0h, 16h, 25h-Rex1high, and 25h-Rex1low) (Kalkan et al., 2017), revealed temporal concordance of the datasets from the two studies (Figure 2A), suggesting that the biological signal driving these temporal clusters is highly reproducible. The transcriptome at 24h post EpiLC induction clustered with those from 16 h and 25h-Rex1high cells (Figure 2A), with the latter previously shown to be in a reversible phase preceding extinction of the naïve state (Kalkan et al., 2017). Consistent with 25h-Rex1high cells, Rex1 (Zfp42) expression in cells at 24 h post EpiLC-induction remained high at the mRNA and protein level (**Tables S2 and S3**). In contrast, the transcriptome at the 36 h time-point during ESC to EpiLC transition clustered with that of the 25h-Rex1low cells, the primary products of exit from naïve pluripotency (Kalkan et al., 2017). Consistent with 25h-Rex1low cells that have exited the naïve ground state, Rex1 expression was downregulated by ∼10-fold at 36 h post EpiLC induction (**Table S3**). Collectively, these data suggest that, by about 36h post induction, cells have exited the naïve pluripotent state.

**Figure 2.**
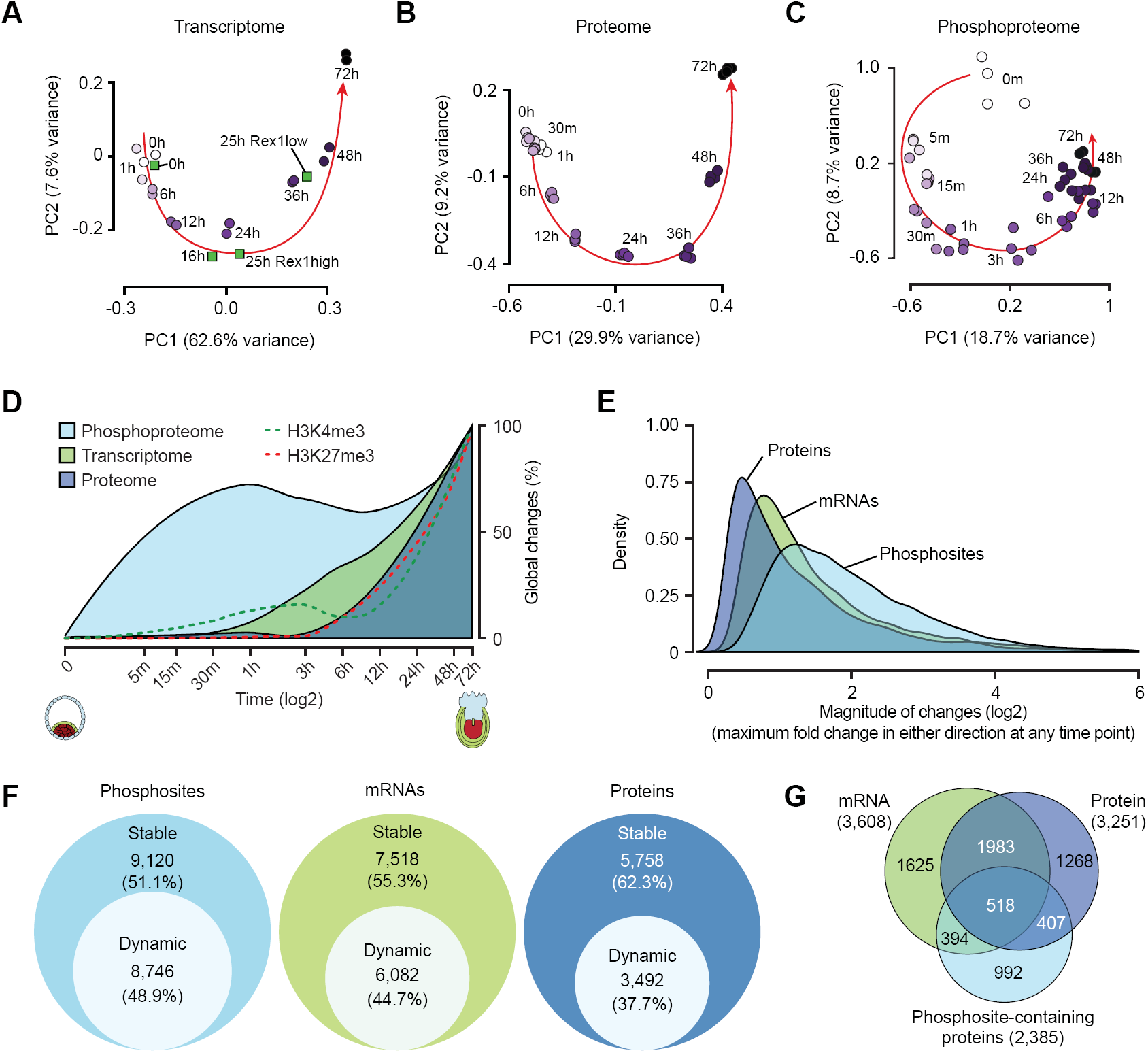
Temporal dynamics of the proteome, phosphoproteome, and transcriptome during ESC to EpiLC transition. (A-C) Principal component analysis (PCA) of the transcriptome (A), proteome (B), and phosphoproteome (C) during EpiLC induction. Each circle represents data from a sample collected at a particular time point during ESC to EpiLC transition, with lighter and darker shades of purple denoting earlier and later time points, respectively. Filled green squares represent transcriptomic data from Kalkan et al. (Kalkan et al., 2017). (D) Temporal dynamics of global changes in the proteome, phosphoproteome, transcriptome, and epigenome during ESC to EpiLC transition. Changes in the phosphorylation level of a given phosphosite were normalized to the changes in corresponding protein level. (E) Density plot showing the distribution of magnitude of changes at the protein, mRNA, and phosphosite level. Changes in phosphosite levels were normalized as in (D). (F) Fraction of phosphosites, mRNAs, and proteins dynamically regulated during EpiLC induction, as assessed using ANOVA test. Changes in phosphosite levels were normalized as in (D). (G) Venn diagram showing overlap among genes encoding differentially regulated mRNAs, proteins, and/or phosphosites during ESC to EpiLC transition. Only genes with both protein and mRNA levels quantified were used for this analysis. See also Figure S3 and S4.

### Phosphoproteome dynamics precede changes to the epigenome, transcriptome, and proteome

To understand the sequence of molecular events and the temporal kinetics that transform cellular identity, we next examined the timing, scale, and magnitude of changes to the proteome, phosphoproteome, transcriptome and epigenome as ESCs transition through various phases of pluripotency. Our analyses revealed that phosphoproteome dynamics precede ordered waves of epigenomic, transcriptomic, and proteomic changes (Figures 2D **and** S4). Notably, about 50% of the regulated phosphosites are significantly modified within 15 min of EpiLC induction, and about one-third were altered as early as 5 min (Figures 2D and S4A). By comparison, <1% of the transcriptome or proteome undergo significant changes within the first hour (Figures S4B **and** S4C). H3K4me3 levels at gene promoters began to change an hour into EpiLC induction, offering the first indication of changes to the epigenome, accompanied by gradual and wide-spread changes to the transcriptome (Figure 2D). While the transcriptome is significantly altered by the sixth hour, widespread changes to the proteome were not evident until about 12 h post-induction (Figures 2D, S4B**, and** S4C), presumably due to latencies associated with protein synthesis and maturation. These data suggest a pioneering role for signaling in pluripotent state transitions.

### Widespread changes to the phosphoproteome mark ESC transition from ground state pluripotency

We next examined the magnitude of changes to dynamically regulated phosphosites, transcripts, and proteins. Our analysis revealed that protein phosphorylation undergoes the greatest degree of change (3.2 median-fold), followed by mRNAs (2.2 median-fold) and proteins (1.8 median-fold) (Figures 2E **and** S4). The broader distribution of the magnitude of changes to the phosphosites (Figure 2E) indicates that the phosphoproteome is more dynamic than the proteome during this transition. Systematic elucidation of differentially regulated phosphosites, mRNAs, and proteins revealed that about half of the phosphoproteome is dynamically regulated over the time-course, whereas only about a third of the proteome undergoes temporal regulation (Figure 2F).

To understand the interplay between protein phosphorylation dynamics and protein abundance, we considered genes whose mRNA, protein, and/or phosphorylation levels were differentially regulated. Our analysis revealed that in ∼28% (925/3,251) of cases, changes at the protein level were associated with significant changes in their phosphorylation level. Notably, among proteins whose abundance was altered, one out of eight (407/3,251) is associated with a significant change to their phosphorylation but not mRNA level (Figure 2G), suggesting a potential role for phosphorylation in regulating the levels and perhaps the activities of a substantial fraction of proteins, presumably by modulating their stability and/or degradation. However, about 60% (1,386/2,385) of the proteins with regulated phosphorylation sites are not associated with significant changes at the protein level, suggesting that phosphorylation/dephosphorylation of these sites may alternatively play a role in altering protein activity, localization, conformation, or interactions. Collectively, these data highlight that changes in the phosphoproteome are rapid, acute, and more widespread than changes in both the transcriptome and proteome, and exemplifies the central role that dynamic phosphorylation plays during the phased progression of pluripotency from the naïve to the primed state.

### *De novo* reconstruction of kinase-substrate networks reveals insights into signaling cascades, dynamics, and crosstalk

To elucidate the set of signaling events, their timing, and order in which they occur as cells transition through various phases of pluripotency, we sought to identify active kinases that underlie signaling cascades. To this end, we used the CLUE algorithm (Yang et al., 2015) to partition all phosphosites into 12 optimal clusters based on their temporal profiles (Figures S5A **and** S5B). Using known kinase-substrate annotations (Hornbeck et al., 2012)., we identified four of these clusters to be enriched for substrates with known kinases: ERK/S6K/RSK, mTOR, p38a, and AKT (Figures 3A, 3B**, and** S5B**; Table S5**). An independent analysis of substrates with known kinases, using our kinase perturbation analysis tool KinasePA (Yang et al., 2016b), confirmed activation/inactivation of these same kinases at various stages during ESC to EpiLC transition (Figure S5C). With the assumption that phosphosites with similar temporal dynamics are more likely to be substrates of the same kinase(s), we hypothesized that proteins containing the phosphosites within each of these four clusters are more likely to be associated with the same signaling pathway. Consistent with this prediction, pathway enrichment analysis of the proteins harboring phosphosites within each of the four clusters revealed enrichment of biological processes strongly associated with the respective kinases (Figure 3C).

**Figure 3.**
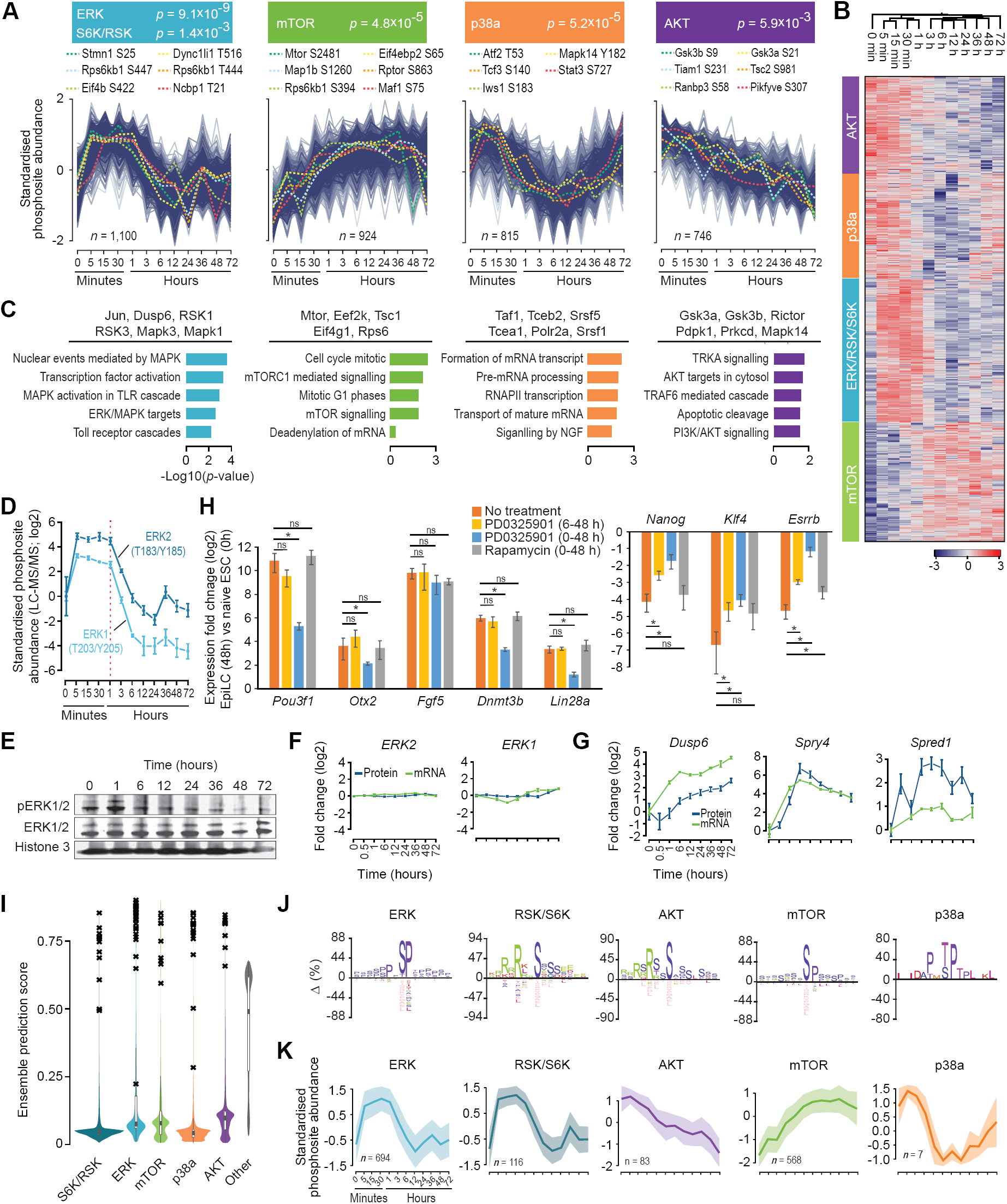
Characterization of signaling dynamics during ESC to EpiLC transition and prediction of substrates for key kinases involved. (A) Clustering of phosphosites based on their temporal dynamics. Four clusters (out of the twelve, see Figure S6A, B) enriched for known substrates of ERK and S6K/RSK (blue), mTOR (green), p38a (orange), or AKT (purple) are shown. Select substrates are highlighted. P-values, Fisher’s exact test. (B) Heatmap representation of the data shown in (A). (C) Gene ontology (GO) analysis of phosphoproteins represented in each of the four clusters in (A). Top five enriched GO categories (biological processes) are shown. Select phosphoproteins within each group are highlighted at the top. (D) Temporal dynamics of relative phosphorylation levels (compared to 0h) of Erk2 (T183/Y185) and Erk1 (T203/Y205, during ESC to EpiLC transition, as quantified using LC-MS/MS. (E) Western blot analysis of total and phosphorylated Erk1/2 during ESC to EpiLC transition. Histone H3 is used as loading control. (F) Temporal dynamics of relative protein and mRNA levels (compared to 0h), as quantified using LC-MS/MS and RNA-Seq respectively, of Erk2 and Erk1 during ESC to EpiLC transition. (G) Same as in (F) but showing data for Dusp6, Spry5, and Spred1, all downstream transcriptional targets of Erk1/2 signaling. (H) RT-qPCR analysis of relative expression of genes associated with naïve pluripotent state (right) or post-implantation epiblasts in EpiLCs (48h) compared to naïve ESCs (0h). During the ESC to EpiLC transition (0-48h), cells were left untreated or cultured in the presence of PD0325901 or Rapamycin for indicated time period. Data, normalized to *Actin*, represents mean of *n* = 3 biological replicates. Error bars represent SEM. *p < 0.05 (Student’s t-test, two-sided). (I) Violin plots showing the distribution of ensemble prediction scores of all profiled phosphosites indicating the likelihood of they being a substrate of one of the five kinases (S6K/RSK, ERK, mTOR, p38a, AKT); kinases other than this five were grouped into the ‘other’ category. Ensemble score for each kinase-substrate pair was generated using a positive-unlabelled ensemble algorithm (Yang et al., 2016a). Black crosses (‘x’) represent previously known substrates. (J) Temporal profiles of predicted substrates for ERK, S6K/RSK, AKT, mTOR, or p38a kinases. Mean and the standard deviation are shown as line-plot and range, respectively. (K) Sequence motifs enriched within predicted substrates for ERK, S6K/RSK, AKT, mTOR, or p38a kinases. Motifs were identified using IceLogo (Colaert et al., 2009), using precompiled mouse Swiss-Prot sequence composition as the reference set. Y-axis represents the difference in the frequency of an amino acid in the experimental vs the reference set. See also Figure S5.

Temporal profiles of the phosphosites within the four clusters revealed the precise timing and order of phosphorylation/dephosphorylation events underlying various signaling cascades (Figure 3B). Notably, substrates within the ERK/S6K/RSK cluster underwent acute phosphorylation within the first five minutes. Interestingly, however, putative ERK substrates, which remained phosphorylated for about an hour after EpiLC induction, reverted to their basal (0h) phosphorylation levels by about 6 h (Figure 3B), suggesting that ERK signaling is inhibited within a few hours after EpiLC induction. Indeed, examination of the phosphorylation dynamics of kinases ERK1 and ERK2 revealed acute dephosphorylation beginning at about an hour after induction (Figure 3D). Consistent with our MS-based phosphoproteomics data, western blot analysis confirmed the transient activation of ERK1/2, with no major changes occurring at the protein or mRNA levels (Figures 3E **and** 3F).

ERK signaling is known to be tightly controlled by negative feedback loops, wherein ERK1/2 activity transcriptionally induces specific ERK1/2 pathway inhibitors, such as dual-specificity MAPK phosphatases (DUSPs), Sprouty (Spry) and Spred proteins, which in turn lead to inhibition and inactivation of ERK1/2 (Caunt and Keyse, 2013; Lake et al., 2016; Ornitz and Itoh, 2015). To assess whether such negative feedback loops shape ERK1/2 signaling dynamics as ESCs transition out of naive pluripotency, we examined the expression dynamics of established negative regulators of ERK1/2 signaling. Within an hour after EpiLC induction, we observed rapid induction of *Dusp6* (an ERK1/2-specific phosphatase), *Spry4*, and *Spred1* (Figure 3G), all downstream transcriptional targets of ERK1/2 signaling (Lake et al., 2016; Lanner and Rossant, 2010). These changes coincided with the acute dephosphorylation of ERK1/2 (Figures 3D **and** 3G), suggesting that ERK1/2 signaling is conceivably under strict control of negative feedback loops as ESCs transition from ground state.

Given the transient activation of ERK1/2 signaling (Figures 3B **and** 3D), we hypothesized that ERK1/2 signaling is perhaps dispensable six hours after EpiLC induction. To test this idea, we added an inhibitor of the MEK/ERK pathway (PD0325901) into the culture medium six hours after EpiLC induction and assessed expression changes of naïve and primed pluripotency factors at 48 h post-induction. ERK1/2 inhibition, beginning at 6 h, had no major effect on the induction of factors associated with primed pluripotency or suppression of naïve pluripotency factors (Figure 3H). However, EpiLC induction in the presence of PD0325901 severely affected both the induction of primed pluripotency factors and the suppression of naïve pluripotency factors (Figure 3H). Altogether, these data establish that while ERK signaling is required to trigger the exit from ground-state naïve pluripotency, it is largely dispensable after about 6 h into EpiLC induction.

Besides ERK1/2, p38 is another MAPK kinase whose known and putative substrates are dephosphorylated within about an hour after EpiLC induction (Figure 3B). Given that ERK1/2-induced DUSP proteins are also known to dephosphorylate the p38 family of MAPKs (Caunt and Keyse, 2013; Lake et al., 2016; Lanner and Rossant, 2010), we examined the temporal dynamics of p38a phosphorylation. Within an hour of ERK1/2 activation, p38a phosphorylation levels decreased by about 4-fold (**Table S1**), suggesting a role for ERK1/2-responsive factors in negatively regulating other pathways, including the p38 MAPK pathway.

ERK1/2-induced Spry and Spred proteins suppress ERK1/2 signaling, in a negative feedback loop, by inhibiting complex formation between the adaptor protein Grb2 and the FGF receptor substrate 2 (Frs2). Intriguingly, the FGF-mediated Grb2-Frs2 signal also regulates the PI3K-AKT pathway as well as other MAPK pathways (p38, JNK) (Lanner and Rossant, 2010; Ornitz and Itoh, 2015). Thus, it is conceivable that any Spry/Spred-mediated negative regulation of Grb2-Frs2 complex formation also inhibits the PI3K-AKT pathway, which is distinct from the MAPK pathways. Indeed, the phosphorylation of known and putative Akt substrates decreased immediately upon ERK1/2 activation (Figures 3A **and** 3B), consistent with pathway cross-talk between ERK1/2 and PI3K-AKT (Mendoza et al., 2011). In addition, withdrawal of LIF to induce EpiLCs (Figure 1A) and subsequent loss of LIF-induced PI3K activation may also have contribute to dephosphorylation of Akt substrates (Yu and Cui, 2016).

Activation of the ERK1/2 pathway, as observed during early stages of ESC transition from the ground state (Figures 3B, 3D**, and** 3E), can also activate the mammalian target of rapamycin complex 1 (mTORC1), an effector molecule downstream of Akt (Mendoza et al., 2011). Active ERK1/2 phosphorylates p90 ribosomal protein S6 kinase (RSK), and together they phosphorylate TSC2 of the TSC complex (which is at the crossroad of ERK1/2 and PI3K-AKT pathways), leading to the release of TSC inhibition of the mTORC1 activity (Mendoza et al., 2011). Consistent with this established link, we find that phosphorylation of known and putative mTOR substrates follows ERK1/2 activation (Figures 3B **and** 3D). To test whether mTORC1 activity is essential for ESC exit from ground state, we induced EpiLCs in the presence or absence of rapamycin, an inhibitor of mTOR that specifically targets mTORC1, and assessed changes in the expression of naïve and primed pluripotency factors at 48 h post-induction. We detected no significant differences (Figure 3H), indicating that mTORC1 activity is not required for exit from naïve pluripotency. Taken together, these data provide insights into extensive cross-talk between signaling pathways and their dynamics during various phases of pluripotency.

### Machine learning predicts substrates for ERK1/2, S6K/RSK, mTOR, AKT, and p38a

Our understanding of signaling pathways that control a vast majority of cell fate decisions is limited because in many cases only a fraction of these pathways has been mapped, with many components remaining to be discovered. Hence, we sought to identify hitherto unknown substrates for each of the key kinases (ERK, S6K/RSK, mTOR, AKT, and p38a) that we had inferred to be active at various time-points during ESC exit from the ground state. We extended our ensemble machine learning algorithm (Yang et al., 2016a) that integrates known kinase recognition motifs and temporal profiles of phosphosites to predict novel substrates for the five kinases of interest (**see Methods**). For each phosphosite and kinase pair, we generated an ensemble prediction score in the range 0 to 1, indicating the likelihood that the phosphosite is a substrate of that kinase. Tabulation of phosphosites by their prediction scores, for each kinase, revealed enrichment of known substrates atop the list (Figure 3I), illustrating the effectiveness of our approach in recovering known substrates.

Using a score cut-off of 0.75, we predicted substrates for ERK, S6K/RSK, AKT, mTOR and p38a kinases (**Table S6**). *De novo* sequence analysis of these substrates identified sequence motifs resembling the consensus recognition motifs of the corresponding kinases (Figure 3J). Despite the similarity between the temporal patterns of predicted substrates for ERK and RSK/S6K (Figure 3K), which are known to act on the same substrate and sometimes in concert (Mendoza et al., 2011), the consensus sequence motifs derived from their putative substrates are quite different. Conversely, although the consensus motifs for predicted ERK and mTOR substrates (or RSK/S6K and AKT substrates) are similar, their temporal patterns are diametrically opposite. These findings illustrate the importance of integrating static features (such as recognition motifs) with dynamic attributes (such as temporal profiles of phosphosites) for successful prediction of novel substrates.

Most signaling cascades culminate in the activation of downstream transcription regulators controlling gene expression programs. Hence, we asked whether transcription regulators, in general, are enriched for dynamically regulated phosphosites. Using the list of annotated transcription regulators (transcription factors, co-factors, and chromatin remodeling enzymes) (Zhang et al., 2012), we found that transcription regulators are more likely to contain dynamically regulated phosphosites than other proteins (odds ratio = 1.91; *p* = 2.1×10^-15^; Fisher’s exact test), suggesting that protein phosphorylation/dephosphorylation could be a general mechanism for modulating the activity of transcription regulators that mediate signal transduction during the pluripotency progression.

To elucidate transcription and chromatin regulators that mediate signaling cascades during the ESC to EpiLC transition, we filtered our list of predicted substrates for known TFs, co-factors, and chromatin modifying enzymes and identified several transcription regulators as putative substrates and possible downstream effectors of ERK, S6K/RSK, mTOR, AKT, or p38a signaling (Figure S5D **and Table S6**). Notably, ERK1/2 is predicted to phosphorylate key transcriptional regulators including Lin28a (RNA binding protein), Zscan4c (expressed transiently in 2-cell embryos and ESCs), EP300 (histone acetyltransferase), Mta3 (member of the Mi2-NuRD histone deacetylase complex) and JunD. Phosphorylation of predicted Lin28a phosphosite (S200) by ERK was recently shown to be an important link between ERK signaling, post-transcriptional gene regulation, and cell fate control (Tsanov et al., 2017). Predicted substrates of mTOR include several chromatin remodeling enzymes with known roles in ESC biology: Jarid2 and Eed (members of the polycomb repressive complex PRC2), Smarca4/Brg1 (the ATPase subunit of the esBAF chromatin remodeling complex), Ino80 (the ATPase subunit of the INO80 chromatin remodeling complex), and Kdm5b (histone H3K4 demethylase). S6K/RSK and AKT are predicted to phosphorylate histone H3K9 demethylase Kdm3b and Dnmt3b, respectively.

### Comparative analysis of changes to the transcriptome and proteome during pluripotency progression

The relationship between mRNA and protein levels is indicative of the combined outcomes of transcription, mRNA stability, translation, and protein degradation (de Sousa Abreu et al., 2009). To understand the downstream effects of signaling on the transcriptome and the extent to which changes at the transcript level during ESC to EpiLC transition translate to changes at the protein level, we examined the temporal dynamics of mRNA expression and protein abundance.

To determine the extent to which mRNA expression captures protein abundance as ESCs transition out of ground-state pluripotency, we assessed the concordance between steady-state mRNA and protein levels at various time-points during EpiLC induction. In agreement with previous studies, which have found a generally limited correlation between steady-state mRNA and protein levels in mammalian systems (Schwanhausser et al., 2011), we found correlation between these layers to be rather moderate and stable across all time-points (Pearson, R = 0.48 to 0.56) (Figure S6A). However, the correlation between changes in mRNA levels and changes in protein levels increased from almost non-existent to moderately high over time (Figures 4A **and** S6B), suggesting that while absolute mRNA levels may not be predictive of protein abundance, changes in transcript level in a perturbed system over a period of time more closely reflect changes in protein abundance.

**Figure 4.**
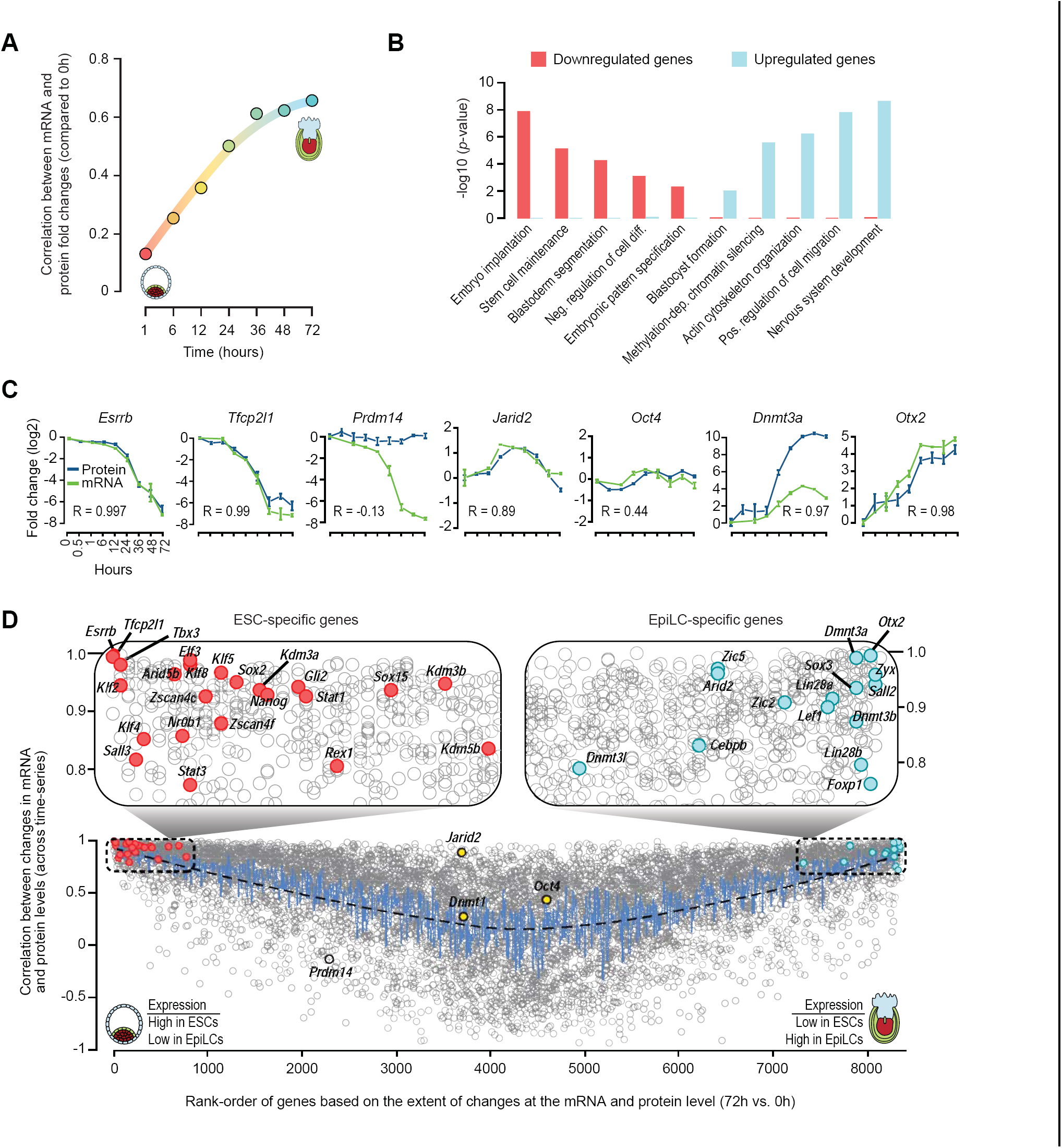
Comparative analysis of the proteome and transcriptome during ESC to EpiLC transition. (A) Temporal dynamics of correlation between changes in protein and mRNA (y-axis) over time in comparison to 0h (ESCs) data. Only genes with both protein and mRNA levels quantified were used for this analysis. (B) Gene ontology (GO) analysis of genes upregulated or downregulated (at both protein and mRNA levels) at 72h vs 0h during ESC to EpiLC transition. Select GO categories (biological processes) enriched among upregulated or downregulated genes are shown. (C) Temporal dynamics of relative protein and mRNA levels (compared to 0h) of select genes. Genes associated with naïve pluripotent state (*Esrrb, Tfcp2l1*, and *Prdm14*), primed pluripotent state (*Dnmt3a* and *Otx2*), and those whose expression is relatively stable during ESC to EpiLC transition (*Jarid2* and *Oct4*) are shown. (D) Correlation between changes in protein and mRNA levels (y-axis) for individual genes plotted against their relative rank-order (x-axis) in terms of change in gene (mRNA/protein) expression (72h vs. 0h) (see Methods). Genes that were substantially down-regulated at 72h vs. 0h have smaller ranks (positioned to the left) and those that were substantially up-regulated have higher ranks (right). Select transcriptional and chromatin regulators, associated with naïve pluripotent state (ESCs), that are downregulated during EpiLC induction are highlighted as filled red circles; those, associated with primed pluripotent state (EpiLCs), which are upregulated during EpiLC induction are highlighted as filled blue circles. Genes whose expression is relatively stable is during EpiLC differentiation are highlighted as filled yellow circles. *Prdm14*, whose protein levels are relatively stable but whose mRNA levels are dramatically downregulated, is highlighted as an open circle. See also Figure S6.

Gene Ontology (GO) analysis of the genes that were down-regulated at both protein and mRNA levels at 72 h vs 0 h revealed enrichment for those associated with stem cell maintenance, blastoderm segmentation, and embryo implantation (Figure 4B). In contrast, up-regulated genes were enriched for those with roles in development and methylation-dependent chromatin silencing (Figure 4B), consistent with significant upregulation of *de novo* DNA methyltransferases Dnmt3a/b/l (Figures 4C, 4D**, and** S6C). Rank-ordering of genes based on the extent of fold-changes at the protein and mRNA level revealed that dynamically regulated genes, including those associated with naïve pluripotent state (e.g., *Esrrb, Tfcp2l1, Nanog, Sox2, Klf2/4, Tbx3, Kdm3a/b*) and post-implantation epiblasts (e.g., *Otx2, Dnmt3a/b, Zic2, Lin28a, Lef1*), exhibit strong correlation (R > 0.85) between changes to the transcript and protein levels (Figures 4C, 4D, S6C**, and** S6D**; Table S7**). Genes whose expression are known to be relatively stable during ESC to EpiLC transition (e.g., *Oct4* and *Dnmt1*) undergo minimal changes and thus exhibit a weak correlation (Figures 4C, 4D **and** S6C). Intriguingly, *Prdm14*, which is expressed in ICM (Yamaji et al., 2008) and downregulated during EpiLC transition (Hayashi et al., 2011; Yamaji et al., 2013), is a notable exception with substantial change at the mRNA but not protein level (Figure 4C**; Table S7**). Consistent with our RNA-Seq and MS-based proteomic data, qRT-PCR and western blot analyses confirmed that while *Prmd14* mRNA level decreases by ∼1,000-fold, its protein level remains unchanged (Figures S1A **and** S6E).

### Distinct waves of global proteomic changes mark discrete phases of pluripotency

Fuzzy *c*-means clustering of the temporal profiles of proteins that were down- and up-regulated during ESC to EpiLC transition revealed a dynamic transposition of cell identity, involving at least three major waves of changes (Figure 5A **and Table S8**). The first wave, presumably induced by upstream signaling events, occurs at about one hour into EpiLC induction and involves down-regulation of naïve pluripotency transcription factors (TFs) including Nanog and Tfcp2l1, both immediate downstream targets of LIF/Stat3 signaling (Niwa et al., 2009), and up-regulation of epiblast-associated factors including Otx2, Zic2, Dnmt3l, and Lin28a. The second wave, at 6 to 24 h, is characterized by downregulation of TFs specific to the naïve pluripotent state and pre-implantation development (Esrrb, Sox2, Tbx3, Nr0b1, and Klf2/4/5), coupled with upregulation of Dnmt3a/b. The third wave coincides with exit from the naïve pluripotent state, at around 36 h, when the cells enter an irreversible phase on their way to establishing a post-implantation EpiLC identity.

**Figure 5.**
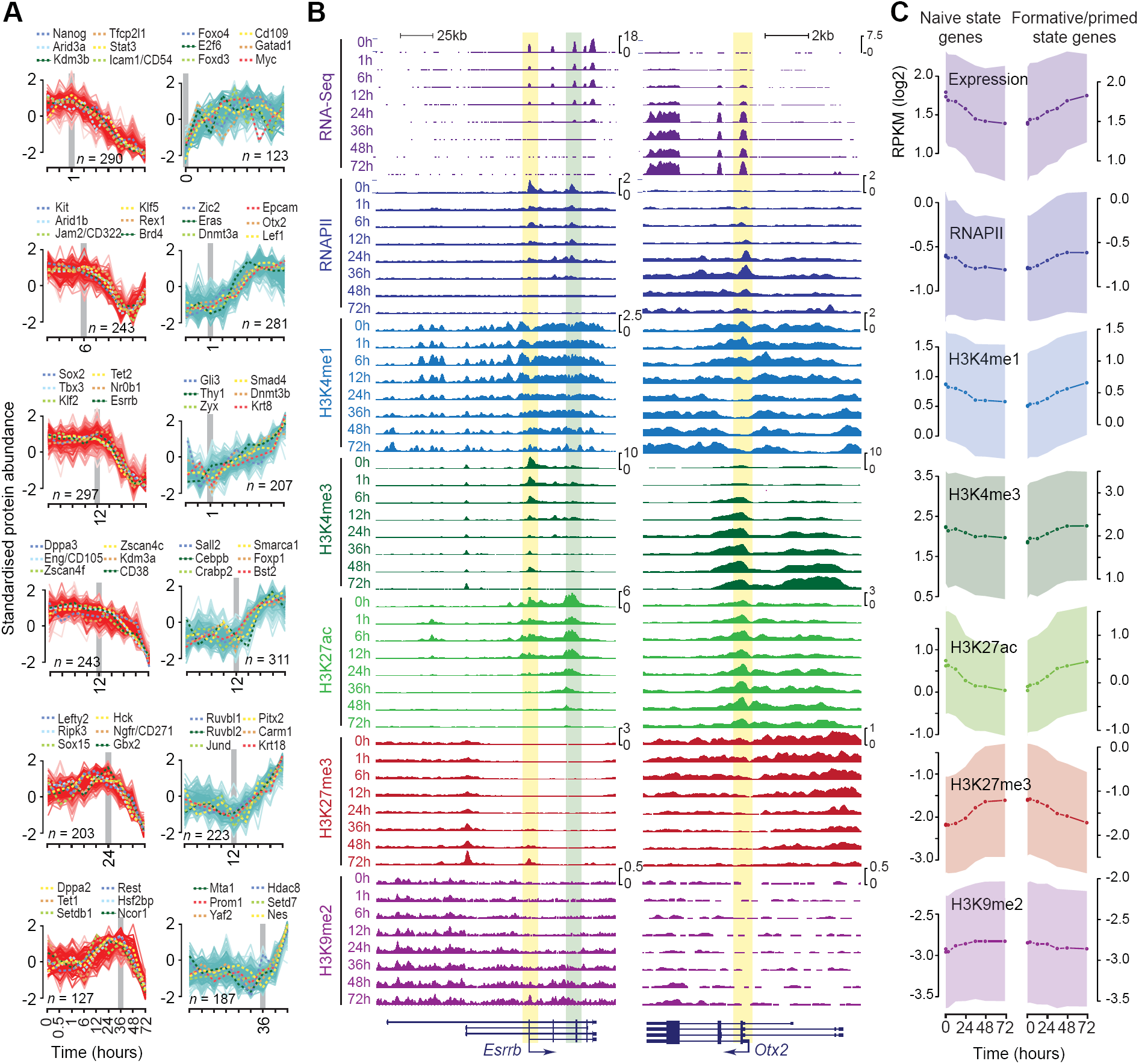
Distinct waves of global changes in the proteome mark various phases of pluripotency. (A) Temporal profiles of standardized changes in protein levels (compared to 0h). Top and bottom 20% of the proteins that are the most down- or up-regulated (red and blue, respectively), based on the rank-ordering in Figure 4D, are grouped into clusters based on fuzzy *c*-means clustering (*c* = 9). Top six clusters, with the most proteins, are shown. Transcriptional and chromatin regulators, known/implicated to play important roles in ESCs and/or EpiLCs, are highlighted. (B) Genome browser shots of *Esrrb* and *Otx2* showing temporal profiles of gene expression dynamics (RNA-Seq) and ChIP-Seq read density profiles for RNAPII and histone modifications H3K4me1, H3K4me3, H3K27ac, H3K27me3, and H3K9me2. Gene annotation is shown at the bottom, with an arrow representing the direction of transcription from the active transcription start site. Regions containing transcriptionally active promoter and known enhancer are highlighted in yellow and green, respectively. (C) Temporal profiles of gene expression, RNAPII, and histone modification dynamics of genes associated with naïve (ESCs) and primed state (EpiLCs). The top and bottom 20% of the genes that are the most down- or up-regulated, based on the rank-ordering in Fig. 4D, were considered as naïve and primed state genes, respectively. Median and standard deviation are shown as line-plot and range, respectively. ChIP-Seq read density within the promoter region was used for analysis (RNAPII, H3K4me3, and H3K27ac: ±1 Kb of TSS; H3K4me1, H3K27me3, and H3K9me2: ±2.5 Kb of TSS).

Chromatin dynamics at the promoters of *Esrrb* and *Otx2* exemplify changes to the epigenome and associated transcriptional output (Figure 5B, C), which precede changes to the proteome. Downregulation of *Esrrb* transcript and transcription-dependent histone modifications (H3K4me3 and H3K27ac) is evident as early as 1 h after EpiLC induction (Figure 5B). Rapid loss of Esrrb transcription is correlated with marked reduction in RNAPII levels at its promoter and is followed by gain of repressive H3K27me3 and H3K9me2 to maintain the repressed transcriptional state. The converse is observed for *Otx2*, wherein loss of H3K27me3 precedes RNAPII recruitment and transcription.

H3K9me2 is known to recruit DNA methyltransferases and is a precursor to DNA methylation (Esteve et al., 2006; Tachibana et al., 2008). Downregulation of H3K9 demethylases (Kdm3a/b) (Figure S6C) coupled with global increase in H3K9me2 levels (Figure 5C) and upregulation of Dnmt3a/b/l (Figures 4C **and** S6C) suggest a finely choreographed sequence of events preceding eventual epigenetic silencing of naïve pluripotency factors by DNA methylation. Together, these data shed light on the tightly orchestrated temporal regulation of gene expression programs that coordinate the transition from naïve to primed pluripotency.

### Identification of cell-surface marker proteins characteristic of various phases of pluripotency

While transgenic reporters can be used to isolate cell populations, cell surface markers allow for prospective identification and tracking of cell types. Given the deep coverage of the quantifiable proteome, we next sought to identify cell-surface proteins characteristic of various phases of pluripotency as ESCs transition from the ground state. Across the profiled time points, we identified 78 cell-surface proteins, representing ∼20% of all cell-surface proteins (Gray et al., 2015), whose expression was quantified with high confidence. Of these 78, 49 were differentially expressed at one or more profiled time points during the ESC to EpiLC transition, of which 34 were at least 3-fold differentially expressed between naïve ESCs (0h) and EpiLCs (72h) (Figure 6A). Most of these cell surface proteins exhibit concordant changes in their transcript levels (Figure S7A), suggesting that changes in their transcript levels accounts for much of the differences in their protein levels. A majority of these cell surface proteins undergo a dramatic transformation in their expression status at around 24-36 h post-EpiLC induction (Figure S7A), presumably coinciding with when the cells exit the naïve pluripotent state to acquire post-implantation epiblast-like identity. These data suggest that the cell-surface proteins captured in our proteomic data set can help discriminate pluripotent cells from pre- and post-implantation epiblast of early mouse embryo.

**Figure 6.**
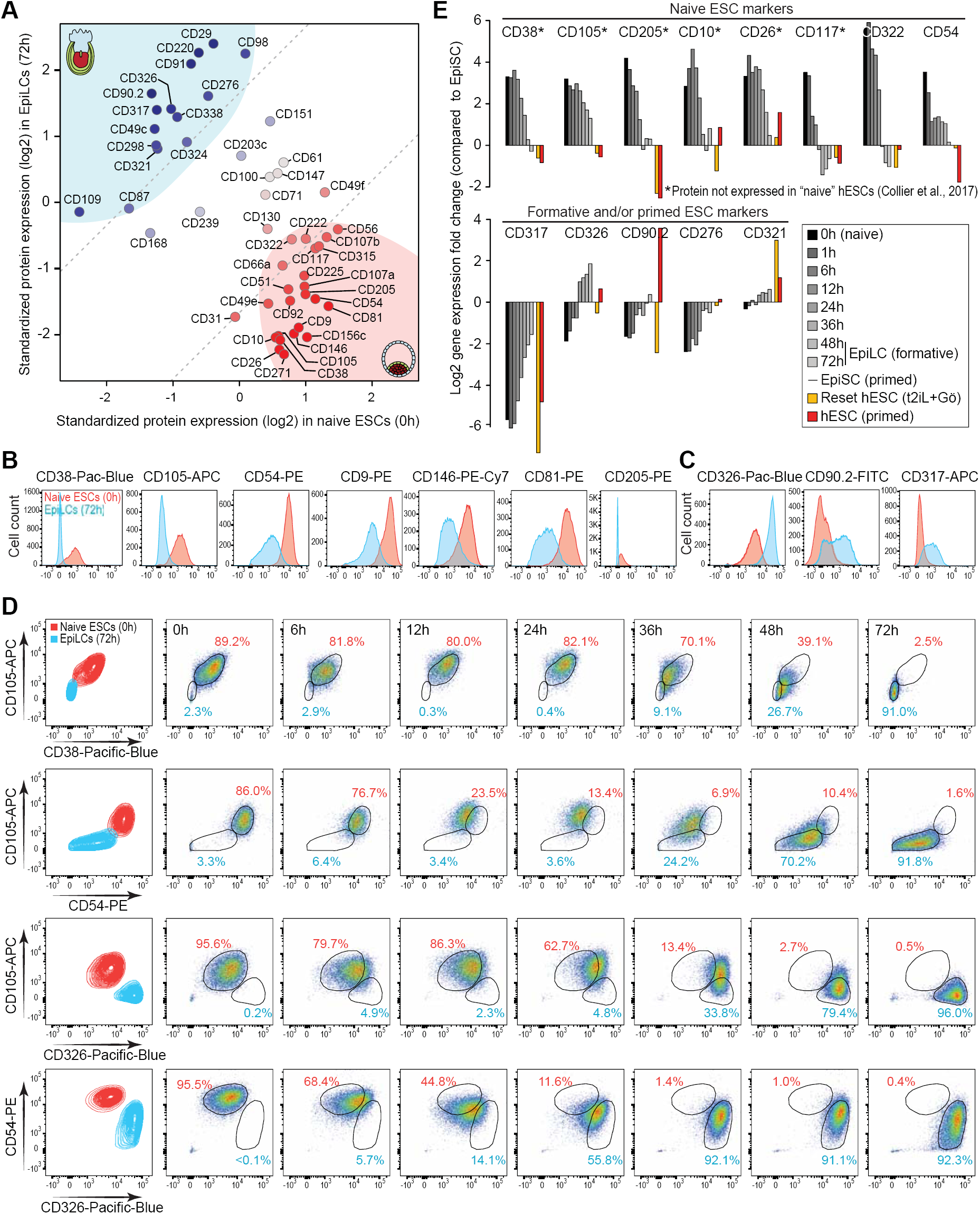
Cell-surface markers specific to naïve and formative/primed pluripotent states. (A) Scatter plot showing expression levels of cell surface proteins in naïve ESCs (x-axis) vs. EpiLCs (y-axis). Data for 49 surface proteins that are differentially expressed at one or more profiled time points during the ESC to EpiLC transition are shown. Based on their distance relative to the diagonal (expressed equally in both cell types), cell surface proteins have been categorized as naïve-specific or primed-specific (darker shades of red and blue, respectively). See Figure S7A for expression dynamics during ESC to EpiLC transition. (B, C) Histograms of flow cytometry analysis using fluorophore-conjugated antibodies showing separation in the fluorescence signal between naïve ESCs (red) and EpiLCs (blue). Data for cell state-specific proteins in naïve ESCs (B) and EpiLCs (C) are shown. (D) Flow cytometry contour plots and dot plots of pairwise antibody combinations in ESCs and EpiLCs (first column) and over the ESC to EpiLC time-course (other columns). (E) Relative gene expression of selected cell surface proteins in mouse and human pluripotent cells based on RNA-Seq data from ESC to EpiLC time-course from this study (0h, 1h, 6h, 12h, 24h, 36h, 48h, and 72h), RNA-Seq data from mouse EpiSCs (Factor et al., 2014), and RNA-Seq data from conventional human ESCs (hESCs) and reset “naïve” hESCs (Takashima et al., 2014). To facilitate direct comparison, all datasets were processed similarly and quantile-normalized. Fold changes relative to expression in mouse EpiSCs are shown. See also Figure S7.

To validate our proteomic data and to define a set of cell surface markers that can discriminate between naïve ESCs and EpiLCs, we performed flow cytometry analysis of candidate cell surface proteins for which antibodies suitable for flow cytometry were commercially available. Our analysis of individual markers with fluorescence-conjugated antibodies revealed a good separation in fluorescence signal between naïve ESCs and EpiLCs (Figures 6B **and** 6C). Consistent with our MS-based proteomic data, CD38 (Adprc1), CD105 (Eng), CD54 (Icam1), CD9, CD146 (Mcam), CD81, and CD205 (Ly75) expression levels are uniformly high in naïve ESCs and low in EpiLCs. Conversely, CD326 (Epcam), CD317 (Bst2), and CD90.2 (Thy1.2) are detected at higher levels in EpiLCs compared to naïve ESCs. Furthermore, flow cytometry analysis of these cell surface proteins during ESC to EpiLC transition revealed that the expression dynamics of individual cell surface proteins faithfully track the phased progression of pluripotency, albeit each protein exhibiting different dynamics during the 72-h time course (Figures S7B **and** S7C). For example, high levels of CD105 and CD38 persisted until 24 h before undergoing downregulation, whereas downregulation of CD54 was more continuous through the time-course. Conversely, while CD326 expression increased gradually over time, upregulation of CD90.2 was not evident until 24 h.

Flow cytometry analysis of multiplexed cell state-specific antibodies showed that combinations of the antibodies can effectively distinguish between naïve ESCs and EpiLCs (Figures 6D **and** S7D). For example, high levels of CD105 and CD38 (or CD54), characteristic of naïve ESCs (Figure 6D), can serve as excellent markers to identify and isolate naïve ESCs from a heterogenous population of pluripotent cells. And, a combination of CD105 (or CD54) and CD326, with discordant expression pattern during ESC to EpiLC transition (Figures S7B **and** S7C), can be useful for tracking the phased progression of pluripotency as ESCs transition from the ground state toward the primed state (Figure 6D). Altogether, these analyses allowed us to identify a robust set of cell state-specific surface proteins, such as CD38, CD105, CD54, CD205, CD10 (Mme), CD26 (Dpp4), CD117 (Kit), and CD322 (Jam2) in naïve ESCs and CD317, CD326, CD90.2, and CD276 in EpiLCs (Figure 6E).

### Comparative analysis of mouse and human pluripotent states

While some cell surface markers specific to “naïve” hESCs, such as CD77 (A4galt) and CD130 (Il6st) (Collier et al., 2017), are also expressed in naïve mouse ESCs (Figure S7E), we found it intriguing that several cell surface markers specific to the naïve mouse ESCs (including CD38, CD105, CD205, CD10, CD26, and CD117) are not expressed in “naïve” hESCs at the mRNA (Figure 6E) or protein levels (Collier et al., 2017). Similarly, CD75 (St6gal1), a marker specifically expressed in “naïve” but not primed hESCs (Collier et al., 2017), is lowly expressed in naïve mouse ESCs compared to EpiLCs or EpiSCs (Figure S7E). Motivated by this lack of concordance, we asked whether the purported naïve hESCs can be considered equivalent to naïve mouse ESCs. If not, this is of interest because they might represent a pluripotent state similar to an intermediate cell state between the naïve and the primed states. To address this question, we compared the transcriptional states of conventional hESCs, considered to be equivalent to mouse EpiSCs (Rossant and Tam, 2017), and hESCs reset to a putatively naïve state (Chan et al., 2013; Takashima et al., 2014) to those of mouse pluripotent cells at various time points during the naïve ESC to EpiLC time-course; as points of reference, we also included data from human blastocyst ICM (Yan et al., 2013), E5.5 mouse post-implantation epiblast (Boroviak et al., 2015), mouse EpiSCs (Factor et al., 2014; Fiorenzano et al., 2016), mouse EpiSCs reset to a naïve-like state (Takashima et al., 2014), EpiLCs (Chen et al., 2018), and conventional mouse ESCs grown in media containing serum+LIF (Fiorenzano et al., 2016; Marks et al., 2012), PD03+LIF (Takashima et al., 2014), or serum+2i+LIF (Chen et al., 2018).

PCA and unsupervised hierarchical clustering revealed distinct clusters of cells corresponding to various pluripotent states (Figures 7A **and** 7B). With much of the variation (35%) captured in the first principal component (PC1), PC1 primarily discriminates between naïve and primed pluripotent states. Conventional hESCs, generally considered as primed (Rossant and Tam, 2017), clustered alongside EpiSCs, considered archetypal representative of primed pluripotency (Rossant and Tam, 2017; Smith, 2017). Interestingly, reset hESCs, reprogrammed to closely resemble mouse naïve ESCs, did not cluster anywhere near naïve mouse ESCs, although they clustered alongside cells from human blastocyst ICM. A closer examination of naïve pluripotency-associated factors in reset hESCs revealed that while the expression of some factors including *Klf4, Klf5, Stella, Prdm14*, and *Zfp42* were reset/upregulated to levels comparable to those in naïve mouse ESCs, many key factors including *Nanog, Esrrb, Nr0b1, Nr5a2, Tfcp2l1*, and *Klf2* were not upregulated to appropriate levels (Figure 7C). Conversely, the expression of many post-implantation epiblast-associated or lineage-specific genes including *Dnmt3a, Dnmt3b, Lin28a, Krt18, Sox4*, and *mir-302b* were not fully downregulated in reset hESCs (Figure 7D). Together, these data suggest that while chemical and/or genetic manipulation of primed hESCs induce molecular features of naive pluripotency in hESCs, reset hESCs are not identical to naïve mouse ESCs.

**Figure 7.**
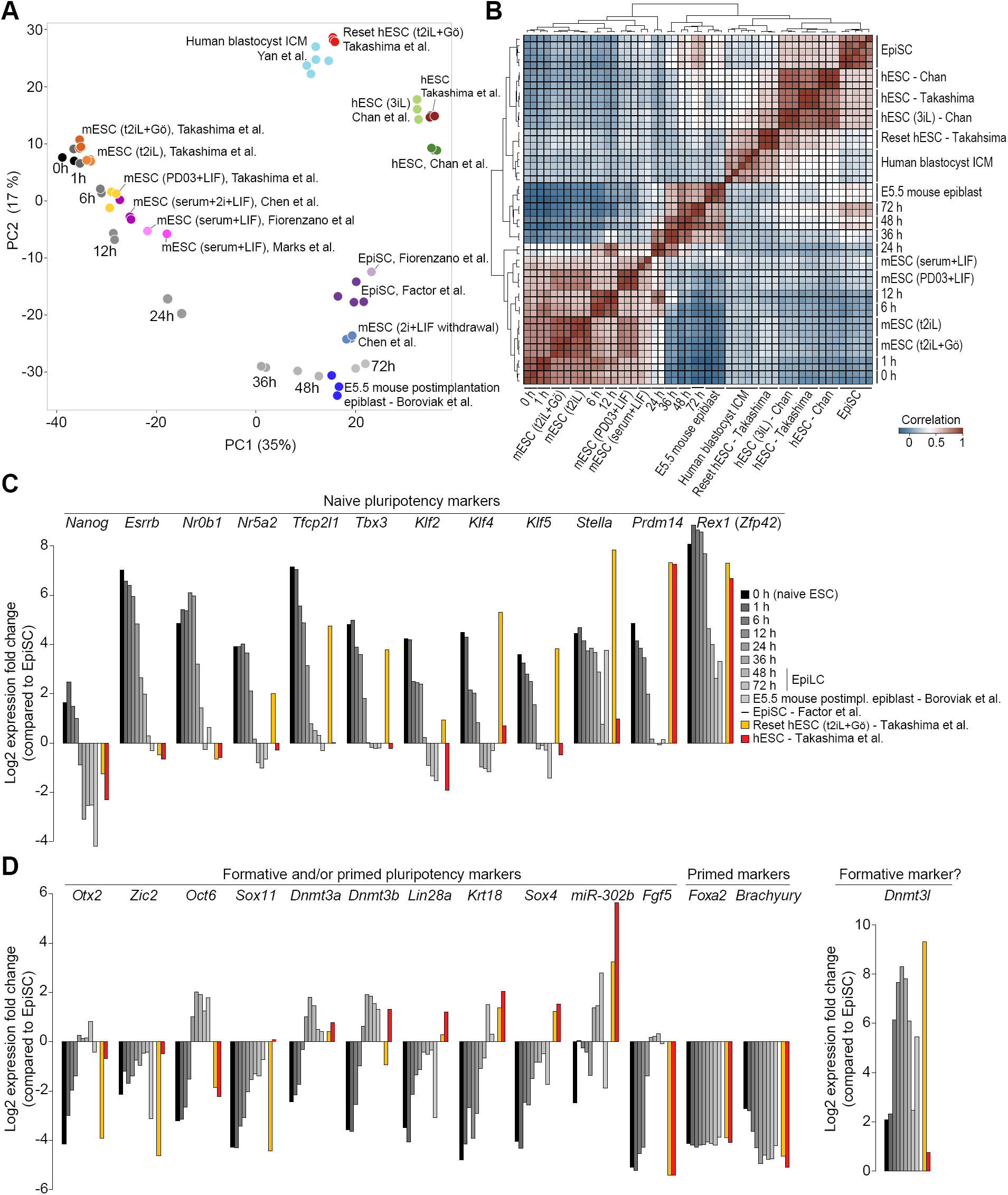
Comparative analysis of mouse and human pluripotent states. (A) PCA of RNA-Seq data from this study (in gray gradient; 0h, 1h, 6h, 12h, 24h, 36h, 48h, and 72h) and previously published studies (in color) (Boroviak et al., 2015; Chan et al., 2013; Chen et al., 2018; Factor et al., 2014; Fiorenzano et al., 2016; Marks et al., 2012; Takashima et al., 2014; Yan et al., 2013). To facilitate direct comparison, all datasets were processed similarly and quantile-normalized. Each data point represents a biological replicate. mESC, mouse ESC; hESC, human ESC. (B) Heatmap showing unsupervised hierarchical clustering of pair-wise Pearson correlations between the RNA-Seq datasets used in (A). (C) Relative expression of genes associated with naïve pluripotency. Fold changes relative to expression in mouse EpiSCs are shown. (D) Same as in (C) but showing genes associated with formative and/or primed pluripotency.

## DISCUSSION

Through integrative analysis of the proteome, phosphoproteome, transcriptome, and epigenome of ESCs transitioning from naïve to primed pluripotency, we have elucidated the sequence of molecular events that underlie the phased progression of pluripotency. Our data data provide new insights into the multi-layered control of developmental transformation from pre-to post-implantation epiblast differentiation and will serve as a rich resource for further investigation of the mechanisms underlying ICM to post-implantation epiblast differentiation.

While previous studies have provided important insights into the proteomes and phosphoproteomes of ESCs in mouse (Christoforou et al., 2016; Li et al., 2011; Nagano et al., 2005; Pines et al., 2011) and human (Brill et al., 2009; Rigbolt et al., 2011; Swaney et al., 2009; Van Hoof et al., 2009), signaling dynamics that underlie pluripotent state transitions remain unexplored. Deeper coverage of the proteome and the phosphoproteome, coupled with high temporal-resolution, allowed us to elucidate signaling dynamics that underlie pluripotent state transitions. Our findings that rapid, acute, and widespread changes to the phosphoproteome precede any changes to the epigenome, transcriptome, and proteome highlight the prominent role signaling plays in cell fate decisions during embryonic development.

*De novo* reconstruction of kinase-substrate networks from our phosphoproteomic data allowed us to elucidate signaling dynamics and extensive crosstalk between signaling pathways during pluripotent state transitions. Consistent with previous studies showing that ERK signaling is required to induce ESCs to a state that is responsive to inductive cues (Kunath et al., 2007), we find that ERK signaling is required to trigger exit from ground-state naïve pluripotency. What was most revealing, however, was the acute dephosphorylation of ERK and its substrates within about six hours into EpiLC induction. This, together with our finding that ERK signaling is largely dispensable after about six hours into EpiLC induction, suggest that the timing and duration of the transient ERK activation is under strict control during pluripotency progression. Indeed, a recent study reported that genetic depletion or chemical inhibition of RSK1, an ERK substrate and a negative regulator of ERK, is sufficient to increase levels of phosphorylated ERK1/2 and alter the kinetics of ESC differentiation (Nett et al., 2018). Conversely, while short-term suppression of ERK signaling helps maintain ESCs in a ICM-like naïve state *in vitro*, prolonged suppression of this pathway compromises the epigenetic and genomic stability as well as the developmental potential of ESCs (Choi et al., 2017).

We also found that mTORC1 activity is not required for exit from naïve pluripotency, consistent with studies showing that mTORC1 activity is not required for cell-fate transition (Betschinger et al., 2013). However, inhibition of both mTORC1 and mTORC2 complexes has previously been shown to induce reversible pausing of mouse blastocyst development and ESCs in culture (Bulut-Karslioglu et al., 2016). Taken together, these findings suggest a requirement for mTORC2 but not mTORC1 for exit from naïve pluripotency.

Our analysis of the phosphoproteome data using a machine learning approach allowed us to predict substrates for key kinases that are active at various phases during pluripotency progression. Our predictions include a number of transcription and chromatin regulators, some of which, we surmise, may have a potential role in mediating/modulating signaling cascades controlling gene expression programs. Further studies are required to validate the predicted substrates, and determine their role, if any, in linking external signals to epigenetic and/or transcriptional programs controlling cell fate transition.

Deep coverage of the proteome, coupled with high temporal-resolution, allowed us to uncover distinct waves of global changes to the proteome that mark discrete phases of pluripotency. The initial wave of changes, likely triggered by the loss of LIF/Stat3 signaling and/or activation of ERK signaling, marks the onset of down-regulation of key naïve pluripotency factors Nanog and Tfcp2l1 along with the activation of post-implantation epiblast markers Otx2 and Zic2. This is immediately followed by the second wave of changes characterized by down-regulation of other naïve markers (Esrrb, Sox2, Tbx3, Nr0b1, Rex1, and Klf2/4/5) and up-regulation of Dnmt3a/b, setting the stage for rewiring of the gene regulatory network and remodeling of the epigenome (Buecker et al., 2014; Kurimoto et al., 2015; Shirane et al., 2016). The final wave of changes, which coincides with the exit from the ground state, likely reflects the completion of the dismantling of the naïve pluripotency network and acquisition of post-implantation epiblast identity. These findings shed the first light on proteome-wide changes during the phased progression of pluripotency.

Because EpiLCs more closely resemble the early post-implantation epiblast (E5.5-E6.5) than do EpiSCs (Hayashi et al., 2011), they have been proposed to represent the “formative” pluripotent state (Rossant and Tam, 2017; Smith, 2017), hypothesized to be the launching pad for multilineage differentiation (Smith, 2017). Although EpiLC induction from ESCs is a directional and progressive process that mirrors epiblast development(Hayashi et al., 2011),The formative state characterized by EpiLCs is transient and cannot be captured in stable self-renewing cell lines using current culture conditions (Hayashi et al., 2011). Our observation that Dnmt3l is transitorily expressed during ESC to EpiLC transition (Figure 7D), coupled with its expression in the epiblast (E4.5-6.5) (Smith et al., 2012) but not in EpiSCs (Veillard et al., 2014), suggests that it could be an excellent marker to isolate formative pluripotent stem cells from a heterogenous population of pluripotent cells. It will be of future interest to determine whether the formative phase can be captured as a stem cell state in culture, as achieved for naïve ESCs and EpiSCs.

Cell surface proteins specific to “naïve” and primed hESCs are known (Collier et al., 2017), but surface markers specific to ground state, as in naïve ESCs, remain to be characterized. Our proteomic data allowed us to identify cell surface proteins that are specific to naïve ESCs and EpiLCs. Flow cytometry analysis using a cohort of antibodies confirmed that the inferred state-specific cell surface markers accurately track pluripotent state transitions, with individual proteins exhibiting different temporal dynamics during the ESC to EpiLC transition. The identified cell surface proteins can enable isolation of specific pluripotent stem cell populations during ESC differentiation and induced pluripotent stem cell (iPSC) reprogramming without having to rely on transgenic reporters.

We were surprised that several cell surface proteins specific to naïve ESCs (CD38, CD105, CD205, CD10, CD26, and CD117) are not expressed in “naïve” hESCs (Collier et al., 2017), raising the question whether the purported naïve hESCs can be considered equivalent to naïve mouse ESCs. The naïve pluripotent state captured in mouse ESCs may be very transient or non-existent in human embryos (Rossant and Tam, 2017). Given the lack of a universal criteria for testing naïve pluripotency in a human system, unlike murine ESCs where chimera contribution to blastocysts is the benchmark, assigning naïve status to reset/reprogrammed hESCs is generally based on molecular but not functional basis (De Los Angeles et al., 2015; Hackett and Surani, 2014). Based on the findings from our comparative analysis of the transcriptional profiles of mouse and human pluripotent states (Figure 7), we propose that the reprogrammed/reset hESCs are more similar to the formative state EpiLCs than to the ground-state naïve mouse ESCs and probably lie somewhere along the developmental axis between the naïve and the formative state.

In summary, our studies provide a comprehensive molecular description of the phased progression of pluripotency. Our data, together with the complementary data describing sequence of molecular events inherent to reprogramming somatic cells into iPSCs (Cacchiarelli et al., 2015; Chronis et al., 2017; Polo et al., 2012; Schwarz et al., 2018), provide a foundation for investigating mechanisms that regulate pluripotent state transitions. The general framework we employed to gain insights into the multi-layered control of pluripotent cell fate transitions is a paradigm that can readily be used to investigate any differentiation process.

## ACKNOWLEDGEMENTS

We thank Gaby Sowa, Igor Paron and Korbinian Mayr for technical assistance with MS measurement. We thank Xiaoling Li, Carmen Williams, and Jothi lab members for critical comments on the manuscript, and Guang Hu for insightful discussion. This work was supported by the Intramural Research Program of the NIH, National Institutes of Environmental Health Sciences (R.J.: 1Z1AES102625) and the Max-Planck Society for the Advancement of Science (M.M.). P.Y. was supported by a Discovery Early Career Researcher Award (DE170100759). mS.J.H. was supported by an EMBO Long-Term Fellowship.

## AUTHOR CONTRIBUTIONS

P.Y., S.C. and R.J. conceived the study; S.C. performed ESC-EpiLC and RNA-Seq experiments; S.J.H. performed phosphoproteomics and proteomics experiments with input from M.M; R.P. performed FACS analyses; A.J.O performed ChIP-Seq experiments. P.Y. performed data analysis with input from S.J.H., D.K., J.Y.H.Y., D.E.J., and R.J.; D.K performed comparative transcriptome analysis of mouse and human cells. P.Y., D.P., and S.J.H developed the data webserver; P.Y., S.J.H., M.M., and R.J. wrote the manuscript; All authors reviewed, edited, and approved the final version of the manuscript.

## SUPPLEMENTAL INFORMATION

**Figure S1. (related to Figure 1).**
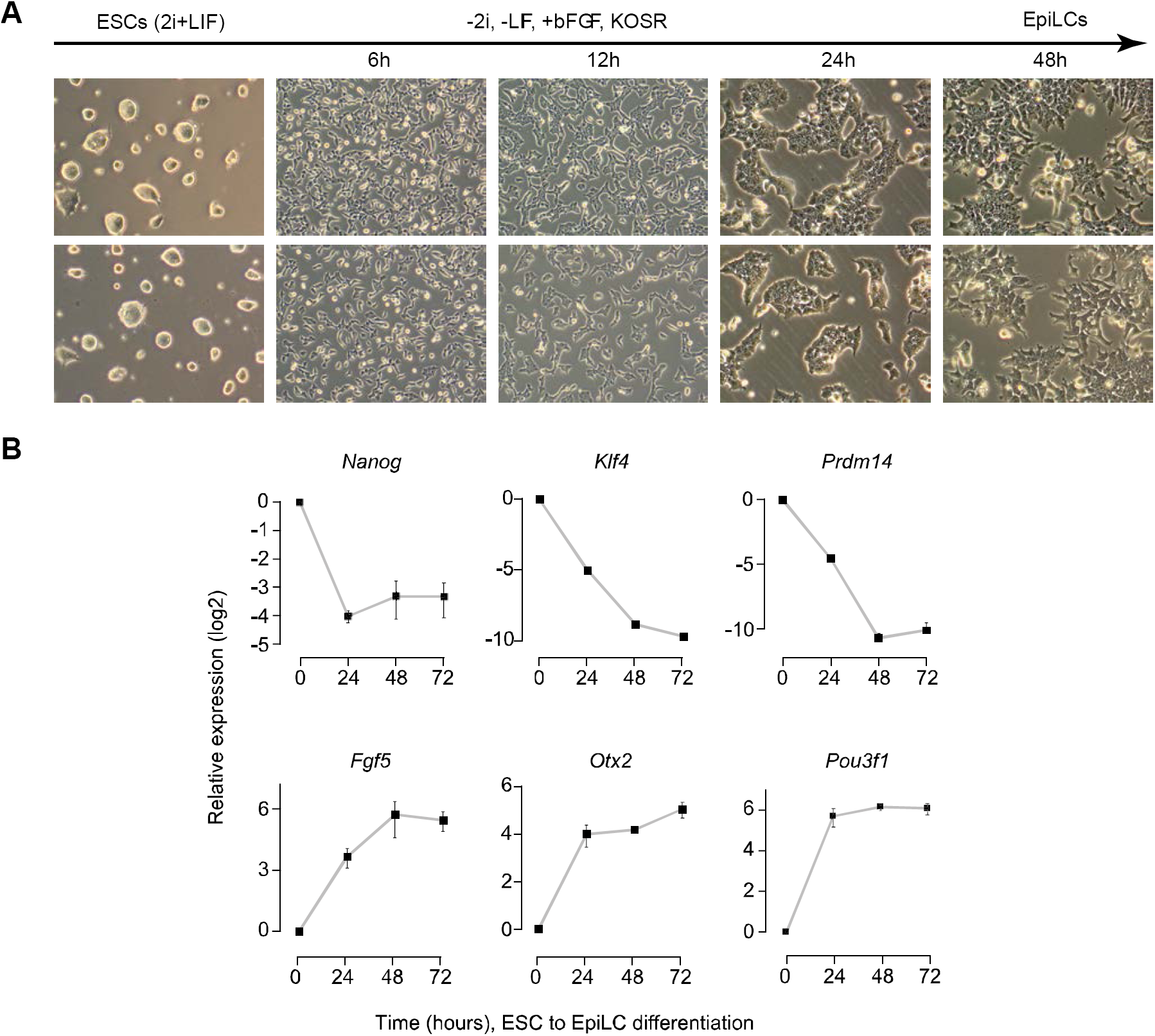
Epiblast-like Cell Induction from ESCs. (A) Morphology of ESCs grown in 2i medium (left-most column; 48h after plating). Other panels show the morphological changes at 6, 12, 24 and 48 hour time-points (after plating) during ESC to epiblast-like cell (EpiLC) transition. Representative images are shown. (B) RT-qPCR analysis of gene expression (mRNA) profiles for pluripotency-associated gene *Nanog*, ICM-associated genes *Klf4* and *Prdm14*, and epiblast genes *Fgf5, Otx2*, and *Pou3f1*/*Oct6* (Buecker et al., 2014; Hayashi et al., 2011) during ESC to EpiLC transition. Data, normalized to *Actin*, represents mean of *n* = 3 biological replicates. Error bars represent SEM.

**Figure S2. (related to Figure 1).**
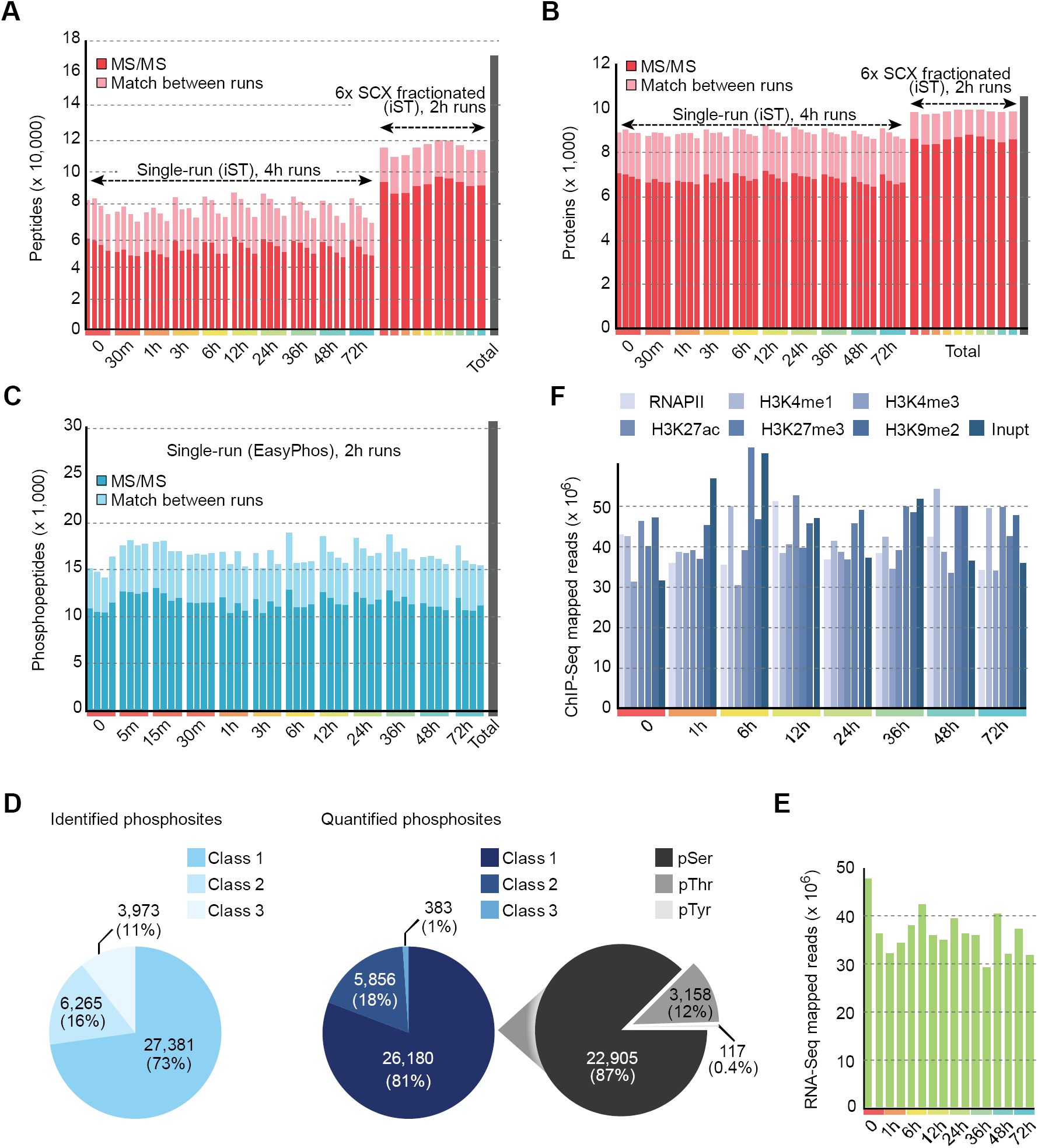
Summary statistics of LC-MS/MS, RNA-Seq, and ChIP-Seq profiling of ESC to EpiLC transition. (A, B) Number of peptides (A) and proteins (B) identified from single-runs and SCX fractionated runs at each profiled time point with/without ‘match between runs’ option in MaxQuant. The union of unique identifications from all time points is represented as total number of peptides/proteins (gray bar). (C) Number of phosphopeptides identified at each profiled time point with/without using ‘match between runs’ option in MaxQuant. The union of unique identifications from all time points is represented as total number of phosphosites (gray bar). (D) Distributions of localization confidence for identified and quantified phosphosites. Classes 1, 2, and 3 represent phosphosites with probability scores >0.75 (high confidence), 0.5-0.75 (medium confidence), and 0.25-0.5 (low confidence) respectively, as quantified by MaxQuant (Tyanova et al., 2016). Phosphorylation types (i.e. pSer, pThr and pTyr) for quantified Class I phosphosites are shown on the right. (E, F) Number of mapped RNA-Seq (E) and ChIP-Seq (F) reads at each profiled time point.

**Figure S3. (related to Figure 2).**
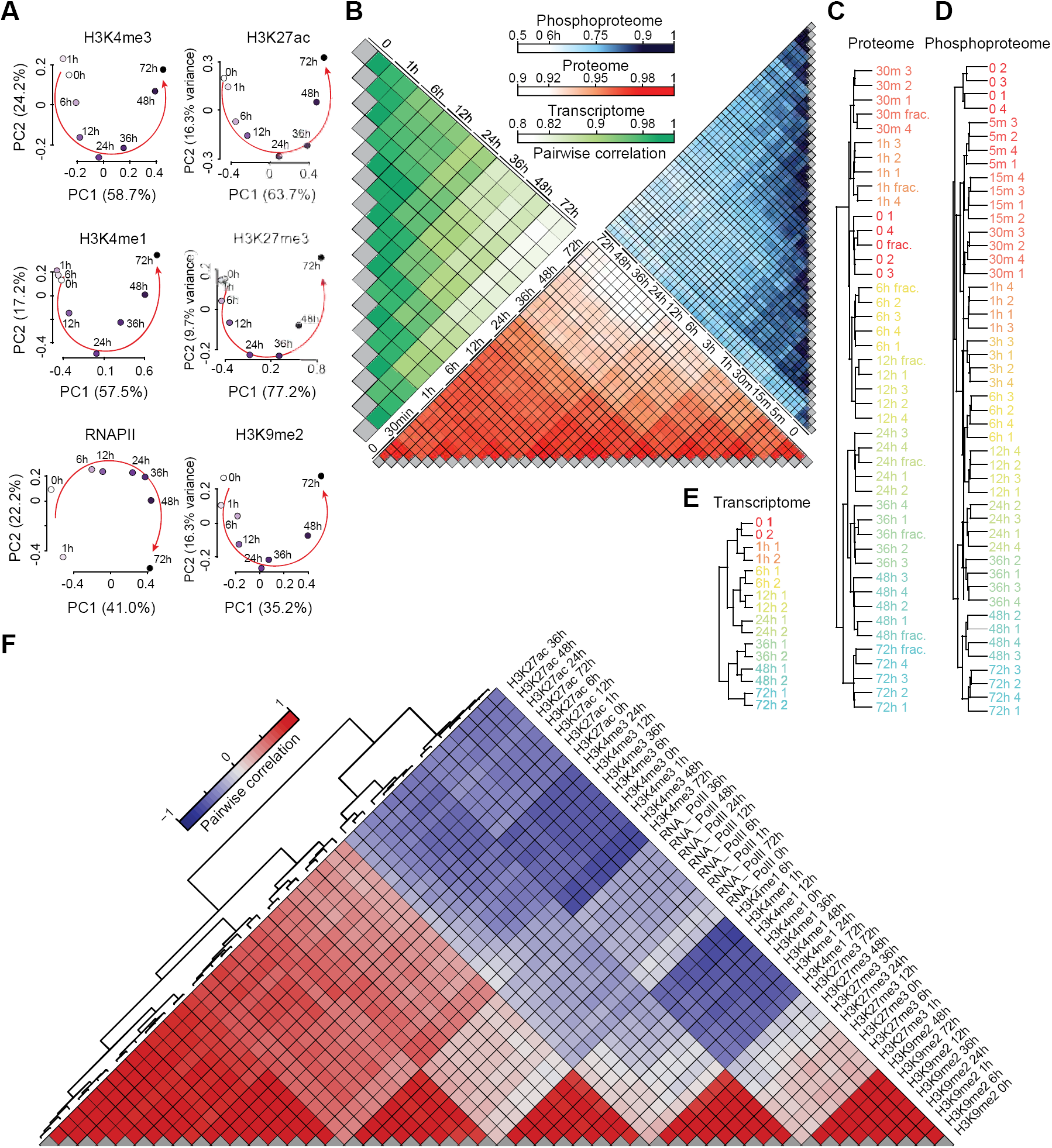
Temporal dynamics of the proteome, phosphoproteome, transcriptome, and epigenome during ESC to EpiLC transition. (A) Principal component analysis (PCA) plot showing temporal dynamics of various histone modifications and RNAPII at gene promoters during ESC to EpiLC transition. (B) Heatmap representation of pairwise correlation between data (transcriptome, green; proteome, red; phosphoproteome, blue) from within (biological replicates) and across all time points. (C-E) Unsupervised hierarchical clustering of proteomic (C), phosphoproteomic (D), and transcriptomic (E) profiles from within (biological replicates) and across all time points. Biological replicates are denoted by suffix 1, 2, 3, or 4. Data from SCX fractionated runs are denoted by suffix “frac.”. (F) Unsupervised hierarchical clustering of pairwise correlation between various histone modifications and RNAPII at gene promoters (±2 Kb of TSS) across all time points.

**Figure S4. (related to Figure 2).**
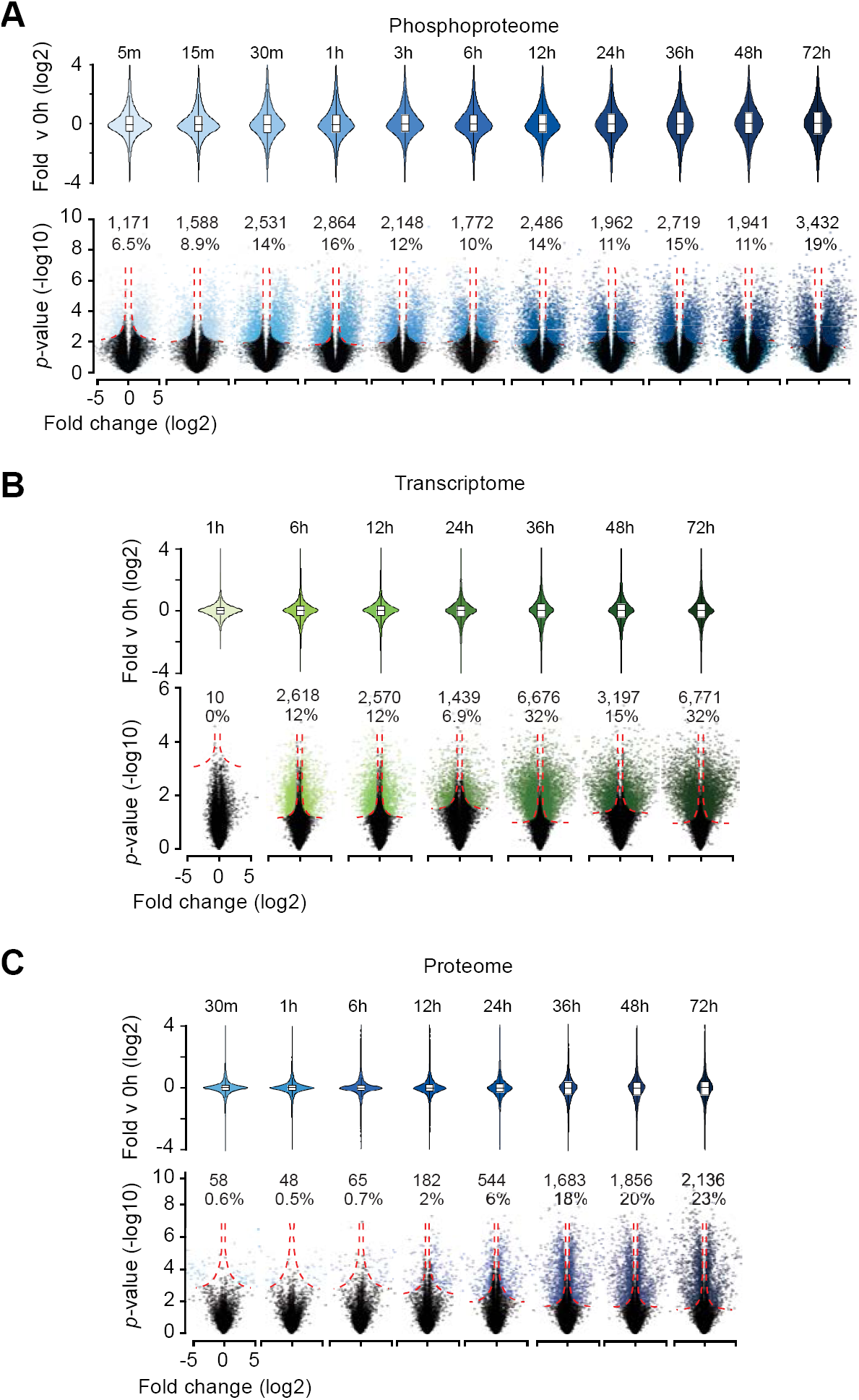
Number and magnitude of differentially regulated phosphosites, genes, and proteins during ESC to EpiLC transition. (A-C) Violin and volcano plots (top and bottom panels, respectively) showing the distribution and magnitude of fold changes for phosphosites (A), mRNA (B), and proteins (C) at each time point in comparison to 0h (ESC) data. Number (and percentage) of phosphosites/mRNAs/proteins that are differentially regulated at each time point (in comparison to 0h), computed using a *t*-test, are shown (refer to Methods for details).

**Figure S5. (related to Figure 3).**
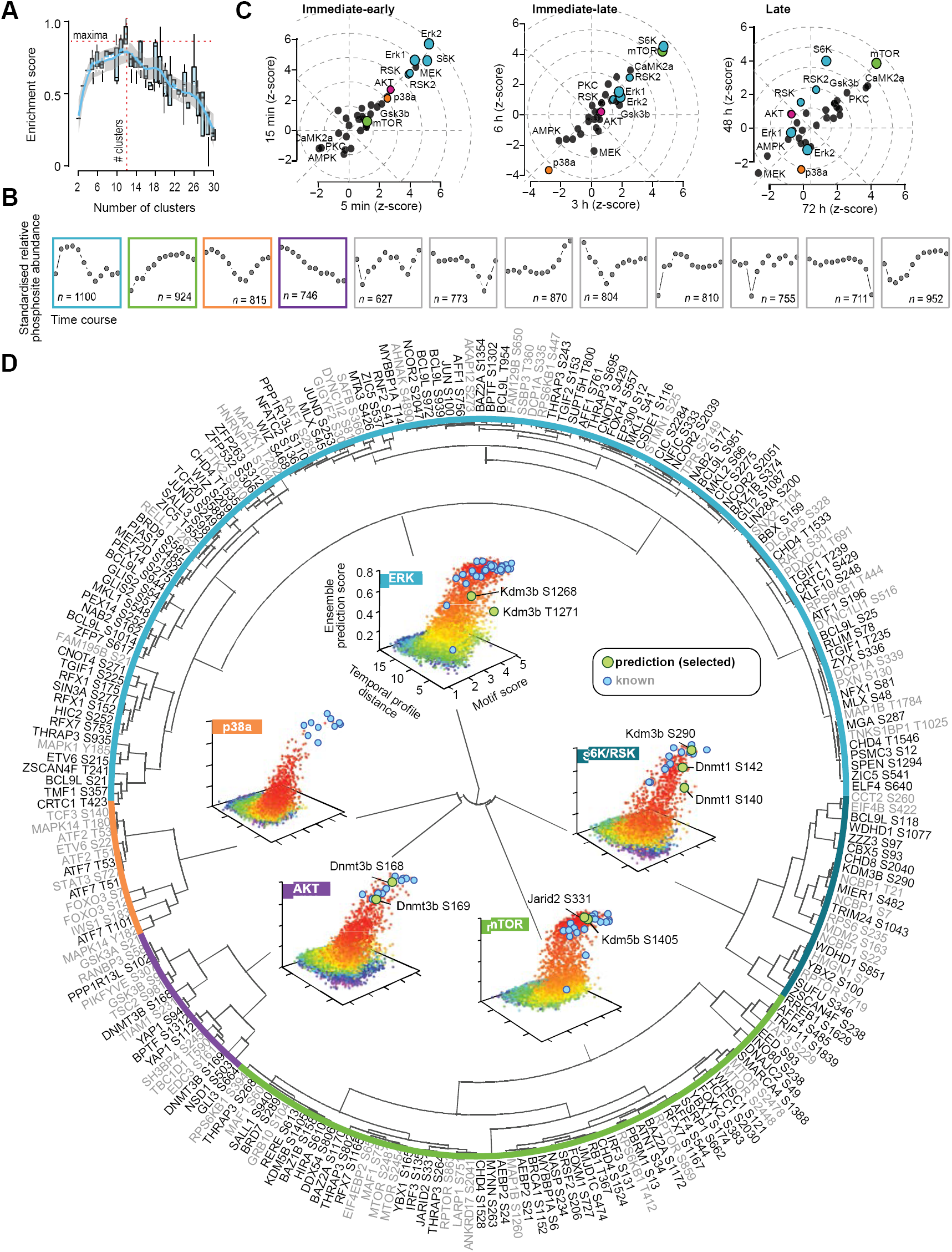
Substrate prediction and characterization. (A) Estimation of the optimal number of clusters of phosphosites using CLUE (Yang et al., 2015). The number of clusters evaluated ranged from 2 to 30, and the optimal number of clusters, as determined by CLUE, was estimated to be 12 based on known kinase-substrate annotations in the PhosphoSitePlus database (Hornbeck et al., 2012). (B) Temporal profiles of standardized, average changes in phosphorylation levels of phosphosites (compared to 0h). Phosphosites are grouped into 12 clusters, as estimated by CLUE. Changes in the phosphorylation level of a given phosphosite were normalized to the changes in corresponding protein levels. Highlighted clusters are enriched for known substrates of AKT (purple), ERK and S6K/RSK (blue), mTOR (green), and p38a (orange). (C) Kinase perturbation plot (Yang et al., 2016b) showing inferred kinase activity (z-score; x-and y-axis) during early (5 and 15 min), intermediate (3 and 6 hours), and late stages (48 and 72 hours) of ESC to EpiLC transition. Z-scores represent relative kinase activity (compared to 0h). (D) Substrate predictions for ERK, S6K/RSK, AKT, mTOR, and p38a kinases. Kinase-substrate prediction using positive-unlabeled ensemble algorithm with multiclass classification (Yang et al., 2016a). Known and predicted substrates within transcriptional and chromatin regulators are shown in gray and black text (on the periphery). See Table S6 for the list of all predictions. Predictions are grouped into clusters based on multiclass prediction scores (refer to Methods for details). Inset: 3D scatter plots showing the ensemble prediction scores (probabilities; z-axis) for each profiled phosphosite with respect to a given kinase (ERK, S6K/RSK, AKT, mTOR, or p38a). X-axis denotes the motif score, representing the enrichment of sequence motif derived from known substrates. Y-axis denotes the dissimilarity (Euclidean distance) between the temporal profile of a given phosphosite and the average temporal profile of known substrates. Color gradient (red to purple) represents high to low ensemble prediction score. Known and select predicted substrates are highlighted in blue and green filled circles, respectively.

**Figure S6. (related to Figure 4).**
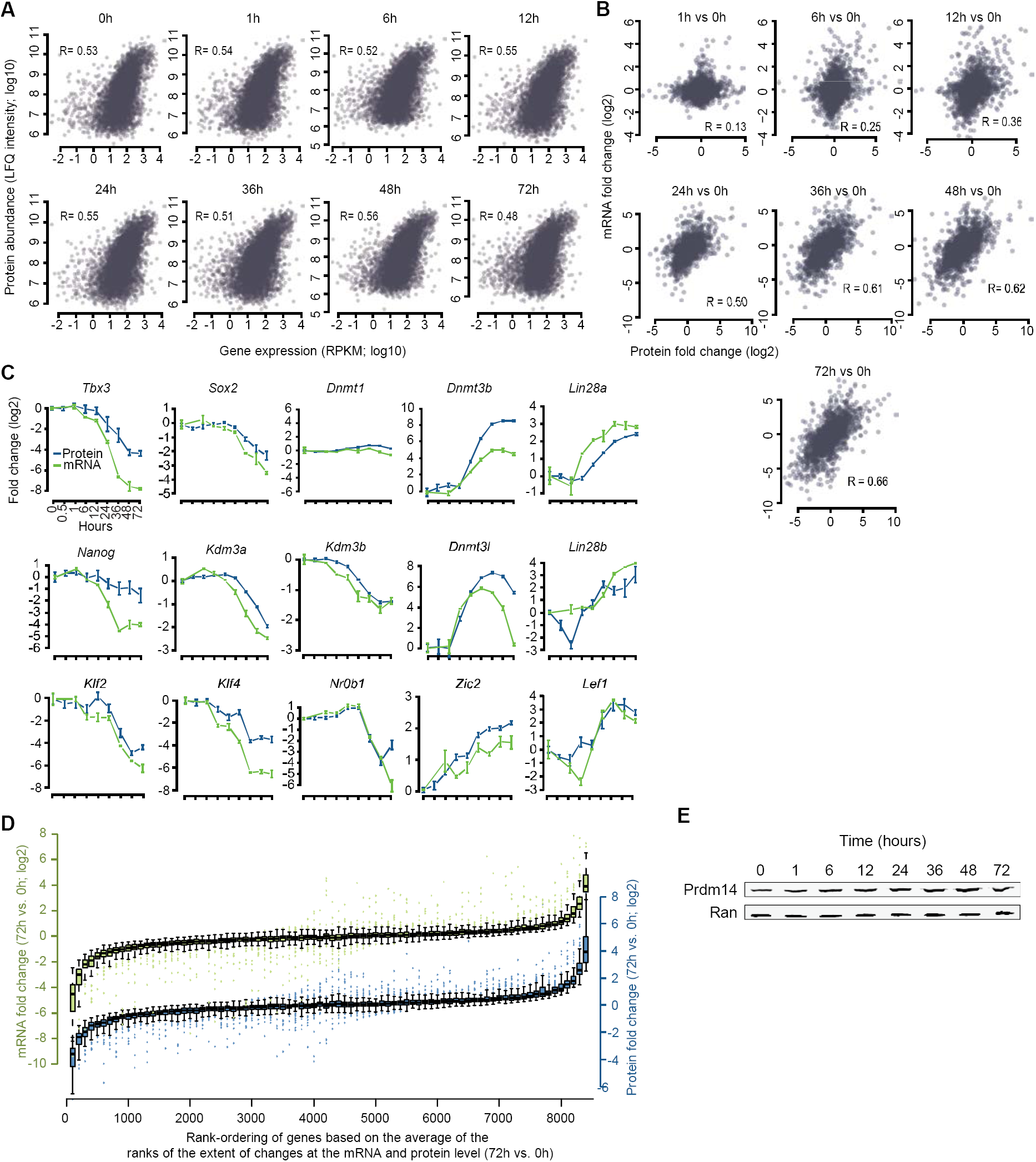
Comparative analysis of the proteome and transcriptome during ESC to EpiLC transition. (A) Scatter plot showing correlation between mRNA (x-axis) and protein (y-axis) levels at each time point. (B) Scatter plot showing correlation between fold-changes in protein (x-axis) and mRNA (y-axis) at each time point in comparison to 0h (ESCs) data. (C) Temporal dynamics of relative protein and mRNA levels (compared to 0h) of select genes. (D) Correlation between changes in mRNA or protein levels (primary and secondary y-axis, respectively) plotted against the relative rank-ordering of genes (x-axis) based on change in gene expression (72h vs. 0h), computed as the average rank of changes in protein and mRNA levels (see Experimental Procedures). (E) Western blot analysis of Prdm14 during ESC to EpiLC transition. Ran is used as loading control.

**Figure S7. (related to Figure 6).**
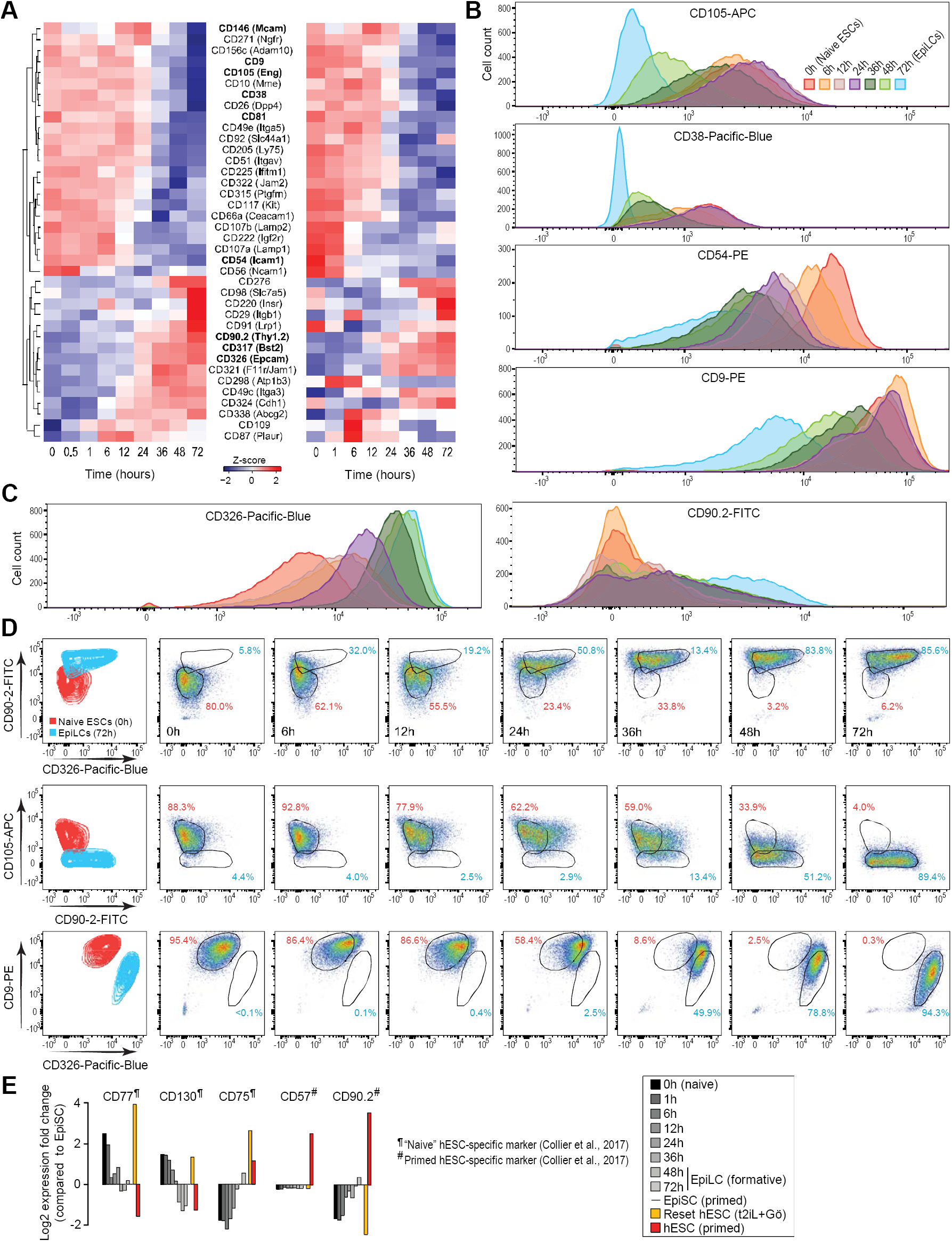
Substrate prediction and characterization. (A) Heatmaps showing temporal profiles of relative protein levels (left) and mRNA expression (right) of cell surface markers during the ESC to EpiLC time-course. Data for 39 markers within the highlighted regions in Figure 6A are shown. Cell surface proteins are ordered based on unsupervised hierarchical clustering. (B) Histograms of flow cytometry analysis using fluorophore-conjugated antibodies against naïve ESC-specific surface markers showing the fluorescence signal tracking the phased progression of pluripotency over the ESC to EpiLC time-course. (C) Same as in (B) but using antibodies against surface markers enriched in EpiLCs compared to naïve ESCs. (D) Flow cytometry contour plots and dot plots of pairwise antibody combinations in ESCs and EpiLCs (first column) and over the ESC to EpiLC time-course (other columns). (E) Relative gene expression of “naïve” and primed hESC-specific cell surface proteins (Collier et al., 2017) in mouse and human pluripotent cells based on RNA-Seq data from ESC to EpiLC time-course from this study (0h, 1h, 6h, 12h, 24h, 36h, 48h, and 72h), RNA-Seq data from mouse EpiSCs (Factor et al., 2014), and RNA-Seq data from conventional human ESCs (hESCs) and reset “naïve” hESCs (Takashima et al., 2014). To facilitate direct comparison, all datasets were processed similarly and quantile-normalized. Fold changes relative to expression in mouse EpiSCs are shown.

## STAR METHODS

Detailed methods are provided in the online version of this paper and include the following:

- KEY RESOURCES TABLE
- CONTACT FOR REAGENT AND RESOURCE SHARING
- EXPERIMENTAL MODEL AND SUBJECT DETAILS
  - Mouse ESC Culture and EpiLC Induction
- METHOD DETAILS
  - Phosphoproteome Sample Preparation
  - Proteome Sample Preparation
  - LC-MS/MS Measurement
  - Quantitative RT-PCR
  - RNA-Seq
  - ChIP-Seq
  - Western Blot
  - Flow Cytometry
  - Phosphoproteomics Data Analysis
  - Proteomics Data Analysis
  - RNA-Seq Data Analysis
  - ChIP-Seq Data Analysis
  - Correlation Analysis of Protein and mRNA expression
  - Comparative Analysis of Multi-ome Dynamics
  - Gene Ontology Analysis
  - Kinase Inference
  - Pathway Enrichment Analysis
  - Substrate Prediction and Motif Analysis
- QUANTIFICATION AND STATISTICAL ANALYSIS
- DATA AND SOFTWARE AVAILABILITY

KEY RESOURCES TABLE.

KEY RESOURCES TABLE
| REAGENT or RESOURCE | SOURCE | IDENTIFIER |
| --- | --- | --- |
| **Antibodies** |  |  |
| Anti-CD105-APC | Biolegend | 120413 |
| Anti-CD146-PE-Cy7 | Biolegend | 134713 |
| Anti-CD205-PE | Biolegend | 359203 |
| Anti-CD317-APC | Biolegend | 127015 |
| Anti-CD326-Pac-Blue | Biolegend | 118225 |
| Anti-CD38-Pac-Blue | Biolegend | 102720 |
| Anti-CD54-PE | Miltenyi Biotech | 130-104-215 |
| Anti-CD81-PE | Biolegend | 104905 |
| Anti-CD9-PE | Biolegend | 124805 |
| Anti-CD90.2-FITC | Biolegend | 105305 |
| Anti-Erk1/2 | Cell Signaling | 9102 |
| Anti-pErk1/2 | Cell Signaling | 9101 |
| Anti-H3K4me1 | Abcam | ab8895 |
| Anti-H3K4me3 | Abcam | ab8580 |
| Anti-H3K9me2 | Abcam | ab1220 |
| Anti-H3K27ac | Abcam | ab4729 |
| Anti-H3K27me3 | Millipore | 07-449 |
| Anti-Histone H3 | Abcam | ab1791 |
| Anti-Prdm14 | EMD Millipore | AB4350 |
| Anti-Ran | BD Bioscience | 610341 |
| Anti-RNA Polymerase II 8WG16 | Covance/Biolegend | MMS-126R |
| IRDye 800CW Goat anti-Mouse IgG | LI-COR Biotechnology | P/N 925-32210 |
| IRDye 800CW Goat anti-Rabbit IgG | LI-COR Biotechnology | P/N 925-32211 |
| IRDye 680CW Goat anti-Mouse IgG | LI-COR Biotechnology | P/N 925-68020 |
| IRDye 680CW Goat anti-Rabbit IgG | LI-COR Biotechnology | P/N 925-68021 |
| **Chemicals, Peptides, and Recombinant Proteins** |  |  |
| 2,2,2-Trifluoroethanol | Sigma | 96924 |
| 2-Chloroacetamide | Sigma | C0267 |
| Acetone | Fisher Scientific | A929 |
| Acetonitrile | Fisher Scientific | A955-4 |
| Accutase | Sigma | A6964 |
| Activin A | Peptrotech | 120-14 |
| Ammonium acetate | Merck | 5438340100 |
| Ammonium bicarbonate | Sigma | 09830 |
| Ammonium hydroxide | Merck | 5330030050 |
| B27 Supplement | Invitrogen | 12587-017 |
| bFGF | Peptrotech | 450-33 |
| BSA fraction V, 7.5% | Invitrogen | 15260-037 |
| CHIR99021, Gsk3 inhibitor | Selleckchem | S2924 |
| DMEM | Thermo Fisher | 11965-084 |
| DMEM/F12 | Invitrogen | 21041-025 |
| DNase I | Worthington | LS002139 |
| DNase I, Amplification grade | Thermo Fisher | 18068015 |
| Dynabeads (protein G) | Pierce | 88847 |
| Dynabeads (protein A) | Pierce | 88846 |
| EDTA | Gibco | 15575-038 |
| EGTA | Bioworld | 40520008-2 |
| FGFR Tyrosine Kinase Inhibitor | Selleckchem | S1470 |
| Fibronectin | Millipore | FC010 |
| Formaldehyde | Sigma | F8775 |
| Formic acid | Fisher Scientific | A11750 |
| Gelatin | Sigma | G1890 |
| Glycerol | Sigma | G5516 |
| Glycine | Sigma | G8898 |
| Guanidine hydrochloride | Sigma | RDD001 |
| GlycoBlue | Ambion | AM9515 |
| HEPES | Sigma | H3375 |
| IGEPAL CA-630 | Sigma | I-3021 |
| KCl | Sigma | P9541 |
| KSR | Invitrogen | 10828-028 |
| Laminin | BD Biosciences | 354232 |
| L-glutamine 200mM | Invitrogen | 25030-081 |
| LiCl | Sigma | 62476 |
| LIF | Millipore | ESG1107 |
| LIVE/DEAD Fixable Near-IR dye | Invitrogen | L34976 |
| LysC | Wako Chemicals | 129-02541 |
| MgCl<sub>2</sub> | Sigma | M2670 |
| N-Lauroylsarcosine | MP Biomedicals | 190110 |
| N<sub>2</sub> | Invitrogen | 17502048 |
| NaCl, 5M | Sigma | S5150 |
| NaHCO<sub>3</sub> | Sigma | S6014 |
| Neurobasal Medium | Invitrogen | 12348-017 |
| NuPAGE Protein Gel | Thermo Fisher | NP0321BOX |
| PBS | Made in house |  |
| PD0325901, MEK inhibitor | S1036 | Selleckchem |
| PMSF | Sigma | P7626 |
| Poly L-ornithine | Sigma-Aldrich | P3655 |
| Protease Inhibitors | Roche | 4693159001 |
| Proteinase K | Invitrogen | 25530049 |
| PVDF membranes | Thermo Fisher | IB24002 |
| QIAzol Lysis Reagent | Qiagen | 79306 |
| RNase cocktail | Ambion | AM2286 |
| SDS, 20% | Fisher Scientific | BP166 |
| Sodium Deoxycholate | Sigma | 30970 |
| SsoFast EvaGreen supermix | Bio-Rad | 1725201 |
| TCEP | Thermo Fisher | PG82080 |
| Trifluoroacetic acid (TFA) | Merck | 8082600100 |
| Tris HCL | Sigma | T2663, T6066 |
| Triton X-100 | Sigma | X100 |
| Trypsin | Sigma | T6567 |
| **Critical Commercial Assays** |  |  |
| iScript cDNA Synthesis Kit | Bio-Rad | 1708891 |
| miRNeasy Mini Kit | Qiagen | 217004 |
| TruSeq stranded total RNA library prep gold | Illumina | 20020598 |
| **Experimental Models: Cell Lines** |  |  |
| Mouse ESCs (E14Tg2a) | ATCC | CRL-1821 |
| **Oligonucleotides** |  |  |
| Primers used for RT-qPCR | This paper | Table S9 |
| **Publicly Available Datasets Used For Analyses** |  |  |
| Mouse ESCs (N2B27+2i) 0h, RNA-Seq | Kalkan et al. (2017) | Array Express: E-MTAB-5305 |
| Mouse ESCs (N2B27) 16h, RNA-Seq | Kalkan et al. (2017) | Array Express: E-MTAB-5305 |
| Mouse ESCs (N2B27) 25h-Rex1high, RNA-Seq | Kalkan et al. (2017) | Array Express: E-MTAB-5305 |
| Mouse ESCs (N2B27), 25h-Rex1low, RNA-Seq | Kalkan et al. (2017) | Array Express: E-MTAB-5305 |
| Mouse ESCs (t2iL), RNA-Seq | Takashima et al. (2014) | Array Express: E-MTAB-2857 |
| Mouse ESCs (t2iL+Gö), RNA-Seq | Takashima et al. (2014) | Array Express: E-MTAB-2857 |
| Mouse ESCs (PD03+LIF), RNA-Seq | Takashima et al. (2014) | Array Express: E-MTAB-2857 |
| Mouse EpiSCs, RNA-Seq | Factor et al. (2014)<br>Fiorenzano et al. (2016) | GEO: GSE57409<br>GEO: GSE79796 |
| Mouse E5.5 postimplantation epiblast, RNA-Seq | Boroviak et al. (2015) | Array Express: E-MTAB-2958 |
| Human ESCs (hESCs), RNA-Seq | Chan et al. (2013)<br>Takashima et al. (2014) | Array Express: E-MTAB-2031<br>Array Express: E-MTAB-2857 |
| Reset hESCs (t2iL+Gö), RNA-Seq | Takashima et al. (2014) | Array Express: E-MTAB-2857 |
| Human ESCs (hESCs, 3iL), RNA-Seq | Chan et al. (2013) | Array Express: E-MTAB-2031 |
| Human blastocyst ICM, RNA-Seq | Yan et al. (2013) | GEO: GSE36552 |
| **Software and Algorithms** |  |  |
| R 3.3.2 | R Core Team, 2016 | https://www.R-project.org/ |
| MaxQuant 1.5.3.29 | Cox and Mann (2008) | http://www.biochem.mpg.de/5111795/maxquant |
| Bowtie 0.12.8 | Langmead et al. (2009) | http://bowtie-bio.sourceforge.net/index.shtml |
| STAR 2.5.2a | Dobin et al. (2013) | https://github.com/alexdobin/STAR |
| Bedtools 2.26.0 | Quinlan (2014) | http://bedtools.readthedocs.io/en/latest/ |
| HTSeq 0.6.1 | Anders et al. (2015) | http://www-huber.embl.de/HTSeq/doc/overview.html |
| ComBat (SVA R package 3.24.0) | Johnson et al. (2007) | https://www.bu.edu/jlab/wp-assets/ComBat/Abstract.html |
| Limma 3.32.2 | Ritchie et al. (2015) | https://bioconductor.org/packages/release/bioc/html/limma.html |
| edgeR 3.16.5 | Robinson et al. (2010) | https://bioconductor.org/packages/release/bioc/html/edgeR.html |
| DESeq2 1.16.1 | Love et al. (2014) | https://bioconductor.org/packages/release/bioc/html/DESeq2.html |
| Clue 1.2 | Yang et al. (2015) | https://CRAN.R-project.org/package=ClueR |
| directPA 1.3 | Yang et al. (2014) | https://CRAN.R-project.org/package=directPA |
| Kinase-substrate Prediction | Yang et al. (2016a) | https://github.com/PengyiYang/KSP-PUEL |
| KinasePA | Yang et al. (2016b) | http://shiny.maths.usyd.edu.au/KinasePA/ |
| IceLogo | Colaert et al. (2009) | https://github.com/compomics/icelogo |

## CONTACT FOR REAGENT AND RESOURCE SHARING

Further information and requests for reagents may be directed to and will be fulfilled by the Lead Contact, Dr. Raja Jothi.

## EXPERIMENTAL MODEL AND SUBJECT DETAILS

### Mouse ESC Culture and EpiLC Induction

Mouse ESCs (E14Tg2a, ATCC) were grown in serum free N2B27-based medium, supplemented with 2i (MEK inhibitor PD0325901, 1.0 μM and Gsk3b inhibitor CHIR99021, 3.0 μM) and LIF (1000u/ml) in tissue culture (TC) plates coated with poly L-ornithine and laminin (Hayashi et al., 2011). For EpiLC induction, ESCs, adapted for a minimum of 4 passages in 2i+LIF, were plated on TC dishes coated with human plasma fibronectin (5µg/ml) in N2B27 medium containing activin A (20 ng/ml), bFGF (12µg/ml) and KSR (1%) (Hayashi et al., 2011).

## METHOD DETAILS

### Phosphoproteome Sample Preparation

All MS experiments were performed in biological quadruplicates. Phosphopeptides were enriched using the EasyPhos workflow as described previously (Humphrey et al., 2015). Briefly, cells were lysed in GdmCl buffer (6M Guanidine hydrochloride, 100 mM Tris pH 8.5, 10 mM TCEP, 40 mM 2-Chloroacetamide) and heated for 5 min at 95°C. Lysates were cooled on ice for 15 minutes, sonicated, and acetone precipitated overnight by addition of 4X volumes of −20°C acetone. Precipitated protein was collected by centrifugation, and pellets washed 1X with 4 Ml −20°C 80% (v/v) acetone. Washed pellets were air-dried for 10 min at room temperature, resuspended in 500 µL TFE digestion buffer (10% TFE (2,2,2-Trifluroethanol), 100 mM ammonium bicarbonate), and sonicated (Bioruptor (Diagenode), 4°C for 2X 5 min cycles) until a homogenous suspension was formed. Protein concentration was determined by BCA assay (Thermo Fisher Scientific). Aliquots corresponding to 1 mg protein were diluted to 500 µL in TFE digestion buffer for phosphopeptide enrichment, and 20 µg protein was used for proteome analysis. Protein was subsequently digested by the addition of 1:100 LysC and Trypsin overnight at 37°C with rapid agitation (2,000 rpm).

### Proteome Sample Preparation

As with phosphoproteome, all MS experiments were performed in biological quadruplicates. In addition, to enhance coverage of the proteome measurements, we pooled the four biological replicates from each time-point and performed StageTip-based Strong Cation Exchange (SCX) fractionation(Wisniewski et al., 2009) of this pooled sample for the proteome runs (Figure 1C, S2A, and S2B). Proteome samples were processed using an in-StageTip (iST) protocol (Kulak et al., 2014), and 10 µg (or 20 µg) protein material was used for single-shot or fractionated samples, respectively. For fractionated samples, equal quantities (5 µg per biological replicate) of protein were pooled prior to digestion. Precipitated protein was reconstituted in iST lysis buffer (6M GdmCl, 100 mM Tris pH 8.5), diluted to 10-fold in iST dilution buffer (10% acetonitrile, 25 mM Tris pH 8.5), and digested with 1:100 LysC (Wako Chemicals) and Trypsin at 37°C overnight directly in StageTips containing SDB-RPS (Styrene Divinyl Benzene Reverse Phase Sulfonate) (3X plugs, Empore 3M) (iST-SDB-RPS) or Strong Cation Exchange (SCX) (6X plugs, Empore 3M) (iST-SCX), for single-shot or fractionated samples respectively. For single-shot iST-SDB-RPS samples, StageTips were washed once with 100 µL 0.2% (v/v) Trifluroacetic acid (TFA), and subsequently eluted with 60 µL 5% (v/v) ammonium hydroxide, 80% (v/v) acetonitrile. For fractionated iST-SCX samples, peptides were eluted in 5X fractions (50 mM, 75 mM, 125 mM, 200 mM, 300 mM) of ammonium acetate, 20% (v/v) Acetonitrile, 0.5% (v/v) formic acid, followed by a final elution with 5% (v/v) ammonium hydroxide/80% (v/v) acetonitrile.

### LC-MS/MS Measurement

Peptides and phosphopeptides were loaded onto a 40 cm column with a 75 μM inner diameter, packed in-house with 1.9 μM C18 ReproSil particles (Dr. Maisch GmbH), and column temperature was maintained at 50°C using a homemade column oven. An EASY-nLC 1000 system (Thermo Fisher Scientific) was interfaced with a Q Exactive HF benchtop Orbitrap mass spectrometer (Thermo Fisher Scientific) using a NanoSpray Flex ion source (Thermo Fisher Scientific). For all samples, peptides were separated with a binary buffer system of 0.1% (v/v) formic acid (buffer A) and 60% (v/v) acetonitrile/0.1% (v/v) formic acid (buffer B), at a flow rate of 300 nL/min. For phosphoproteome analysis peptides were eluted with a gradient of 5% - 25% buffer B over 85 minutes followed by 25% - 55% buffer B over 45 minutes, and peptides were analysed with one full scan (300-1,600 m/z; R=60,000 at 200 m/z) at a target of 3e6 ions, followed by up to five data-dependent MS/MS scans with HCD (target 1e5 ions; max IT 120 ms; isolation window 1.6 m/z; NCE 25%; 40% underfill ratio), detected in the Orbitrap detector (R=15,000 at 200 m/z). Dynamic exclusion (40 s) and Apex trigger (4 to 7 s) were switched on. For single-run proteome analysis, peptides were eluted with a gradient of 4% - 32% buffer B over 180 minutes followed by 32% - 47% buffer B over 40 minutes, and for pooled SCX-fractionated samples, peptides were eluted with a gradient of 4% - 32% buffer B over 90 minutes followed by 32% - 47% buffer B over 20 minutes. Peptides were analysed, with one full scan (300-1,600 m/z; R=60,000 at 200 m/z) at a target of 3e6 ions, followed by up to 10 (for single-run samples) or 15 (for fractionated samples) data-dependent MS/MS scans with HCD (target 1e5 ions; max IT 100 ms for single-run samples, 25 ms for fractionated samples; isolation window 1.6 m/z; NCE 25%; 30% underfill ratio), detected in the Orbitrap detector (R=15,000 at 200 m/z). Dynamic exclusion (30 s) was switched on.

### Quantitative RT-PCR

Quantitative RT-PCR was performed as previously described (Oldfield et al., 2014). Briefly, Total RNAs were prepared from cells using Qiazol lysis reagent (Qiagen), and cDNAs were generated using the iScript kit (Bio-Rad) according to the manufacturer’s instructions. Quantitative PCRs were performed on the Bio-rad CFX-96 or CFX-384 Real-Time PCR System using the Bio-rad SsoFast EvaGreen supermix. Three or more biological replicates were performed for each experiment. Data are normalized to *Actin* expression, and plotted as mean +/-S.E.M. See Supplementary Table 9 for primers used in RT-qPCR analysis.

### RNA-Seq

Total RNA was extracted with Qiazol lysis reagent (Qiagen) treatment and purified using miRNeasy Kit. The samples were then treated with DNase I, Amplification grade (Invitrogen) and stranded libraries were prepared using the TruSeq stranded RNA kit (Illumina) with RiboZero depletion (Gold kit) and sequenced on Illumina HiSeq system.

### ChIP-Seq

ChIP was performed as previously described (Oldfield et al., 2014). Briefly, mouse ESCs (1×10^7^) were cross-linked with 1% formaldehyde in DMEM for 10 min, and the reaction was quenched by the addition of glycine at a final concentration of 125 mM for 5 min. Cells were washed twice with PBS, and resuspended in 1 ml of lysis buffer A (50 mM HEPES pH 7.5; 140 mM NaCl; 1 mM EDTA; 10% Glycerol; 0.5% IGEPAL CA-630; 0.25% Triton X-100; 1x Complete protease inhibitor mixture, 200 nM PMSF). After 10 min on ice, the cells were pelleted and resuspended in 1 ml of lysis buffer B (10 mM Tris-HCl pH 8.0; 200 mM NaCl; 1 mM EDTA; 0.5 mM EGTA; 1x protease inhibitors, 200 nM PMSF). After 10 min at room temperature, cells were sonicated in lysis buffer C (10 mM Tris-HCl pH 8.0; 100 mM NaCl; 1 mM EDTA; 0.5 mM EGTA; 0.1% sodium deoxycholate; 0.5% N-lauroylsarcosine; 1x protease inhibitors, 200 nM PMSF) using Diagenode Bioruptor for 16 cycles (30 sec ON; 50 sec OFF) to obtain ∼200–500 bp fragments. Cell debris were pre-cleared by centrifugation at 14,000 rpm for 20 min, and 8 μg (or 20 μg) of chromatin was incubated with antibodies against specific Histone modifications (or RNA Pol II, respectively) overnight at 4 °C. Protein A/G-conjugated magnetic beads (Pierce Biotech) were added the next day for 2 hours. Subsequent washing and reverse cross-linking were performed as previously described (Heard et al., 2001).

### Western Blot

Western-blots were performed as previously described (Oldfield et al., 2014). Briefly, Cell pellets, lysed in RIPA buffer (25 mM Tris-HCl, pH 7.4, 150 mM NaCl, 1% IGEPAL, 1% Sodium deoxycholate) with protease inhibitors, were sonicated using Bioruptor (Diagenode) for three cycles (30 sec ON; 50 sec OFF). The lysate was boiled with SDS-PAGE sample buffer, loaded onto NuPAGE gel, and transferred to 0.22 μM PVDF membranes. The membranes were pre-wet in 100% methanol and rinsed with ultrapure water before being washed for 5 min in 1x PBS. The membranes were then blocked with Odyssey blocking buffer for 1 h at room temperature with gentle shaking. Each membrane was treated with appropriate primary and secondary (IRDye) antibodies. The membranes were then washed in PBS (0.1% Tween 20), rinsed with PBS and scanned and quantified on an Odyssey imaging system.

### Flow Cytometry

Cells were dissociated into single cells with Accutase, washed and passed through 40 μm cell strainers. Cells were washed with PBS and stained with LIVE/DEAD Fixable Near-IR dye (Invitrogen) to stain dead cells (1 x 10^6^ to 2 x 10^6^ cells per reaction). Cells were washed 2X with flow buffer (2% FBS in PBS, 1Mm EDTA, 25ug/ml Dnase I). Conjugated antibodies were mixed with 50 μL flow buffer and applied to 50 μL of cells. Cells were incubated for 30 minutes at 4°C in the dark and washed 2X with buffer (2% FBS in PBS) and centrifuged at 300xg for 5 minutes. Data was analyzed using FlowJo V10 software or FACSDiva (BD Biosciences).

### Phosphoproteomics Data Analysis

Raw MS files from phosphoproteomics experiments were processed using MaxQuant (version 1.5.3.29) (Cox and Mann, 2008) for phosphosite identification using mouse UniProt (August 2015 release). In total, 37,619 phosphorylation sites were identified, which are classified into Class I (27,381), II (6,265) and III (3,973) based on MaxQuant reported confidence of localization scores (Figure S2A, left panel). Phosphorylation level of each site was quantified using LFQ intensity from MS and logarithm (base 2) transformed. Denoting the 12 profiled time points as *t*_*i*_ (*i* = 0, 5m, 15m, 30m, 1h, 3h, 6h, 12h, 24h, 48h, 72h) and the number of times a phosphorylation sites (*p*) quantified at a given time point as *q*^*p*^(*t*_*i*_). Phosphorylation sites from Class I were filtered to require at least 4 valid values in any one of the 12 time-points (i.e. ∃ *i* such that *q*^*p*^(*t*_*i*_) = 4 Subsequently, only phosphorylation sites with at least 12 out of 48 quantified values (12 time-points, four replicates) were retained (i.e. ∑ _*i*_ *q*^*p*^(*t*_*i*_) ≥ 12 for further analysis. This resulted in 17,866 phosphorylation sites passing the above stringent filtering criteria. This filtered data was then median-normalized with respect to each of the 12 time-points and remaining missing quantifications within these data were subsequently imputed using a two-step procedure. In the first step, for each phosphorylation site with two or more quantified values out of the four biological replicates in each time point (i.e. ∃ *i* such that *q*^*p*^(*t*_*i*_) ≥ 2, we calculated the mean 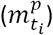and standard deviation 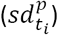 for that *p* at *t*_*i*_ using quantified replicates and imputed missing data for *p* at *t*_*i*_ using a Gaussian model parameterised by 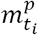 and 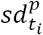. In the second step, we imputed the remaining missing values using the heuristic random-tail method described previously (Robles et al., 2017). Specifically, for each time point *t*_*i*_ the grand mean 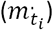 and grand standard deviation 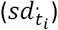 across all phosphorylation sites were calculated and a Gaussian model were utilised to impute missing data in each *t*_*i*_ by down-shifting 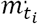. by 1.6 and with a standard deviation of 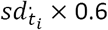 Phosphoproteomics data were subsequently corrected for batch effects using ComBat (Johnson et al., 2007), and finally data was normalized by the total proteome.

### Proteomics Data Analysis

Like the phosphoproteome data, raw MS files from total proteome experiments were processed using MaxQuant (1.5.3.29) for protein identification using mouse UniProt database (August 2015 release). After filtering to remove common protein contaminants and reverse matches, we identified a total of 10,597 proteins. Protein abundance was quantified using LFQ intensity and log (base 2) transformed. Since the fractionated samples have fewer missing values (Figures S2B and S2C), we took advantage of the more complete quantitation from fractionated samples to guide the imputation of missing values in the single-run samples. Denoting the 9 profiled time points in proteomics experiment as *t*_*i*_ (*i* = 0, 30m, 1h, 3h, 6h, 12h, 24h, 48h, 72h) and a protein (*p*) that is quantified at a given time point in the fractionated sample as *s*^*p*^ (*t*_*i*_), protein identifications from fractionated samples were filtered to require at least 5 valid values out of the 9 time-points (i.e. ∑_*i*_*s*^*p*^ (*t*_*i*_ ≥ 5). Then, for fractionated samples we calculated the means 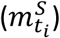 and the standard deviations 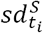 at each time point *t*_*i*_ and imputed the missing values in the fractionated samples at each time point by downshifting 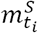 by 1.8 and with a standard deviation of 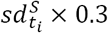 as in Beck et al., 2015. After filtering and imputing data specifically for fractionated samples, we first calibrated the single-run samples with respect to fractionated samples at each time point and then imputed missing values by using the means 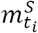 and standard deviations 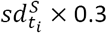 calculated from fractionated samples.Then, batch effect correction was applied using ComBat (Johnson et al. 2007 Biostatistics) for subsequent analysis. Fuzzy *c*-means clustering (*c* = 9) was used to partition the proteins that are the most down-regulated or up-regulated into clusters based on their temporal expression profiles (Figure 5A). Resulting clusters were ranked by the cluster size (number of proteins) from large to small, and the top six clusters, with the most proteins, are shown (Figure 5A).

### RNA-Seq Data Analysis

Pair-end 51 bp reads were mapped to the mouse (mm9) genome using STAR (version 2.5.2a) (Dobin et al., 2013), allowing up to three mismatches, retaining only reads that align to unique locations, and permitting a maximum intron length of 100,000. For visualization on the UCSC Genome Browser and generation of screenshots, mapped reads were normalized to reads per million (RPM) and plotted as histograms using Bedtools version 2.26.0 (Quinlan, 2014). For gene expression analysis, mapped reads were subsequently used to quantify Ensembl/Refseq transcript and gene models (Flicek et al., 2012) using HTSeq version 0.6.1 (Anders et al., 2015). Raw read counts per gene were normalized using the DESeq2 R package version 1.16.1 (Love et al., 2014), batch effect corrected by ComBat, and transformed using a regularized log function implemented in DESeq2. Gene length was extracted from BioMart Database (Durinck et al., 2005), and edgeR package version 3.18.1 (Robinson et al., 2010) was used to calculate RPKM for each gene. RNA-Seq data from Kalkan et al. (2017) were processed similarly (as described above) and normalized together with RNA-Seq data generated for this study using DESeq2 to facilitate principal component analysis (PCA) (Figure 2A). For comparison of RNA-Seq data from mouse and human cells (Figures 6E, 7C, and 7D), RPKM data for genes were log2 transformed (after adding 1) and quantile normalized. For PCA and unsupervised hierarchical clustering of RNA-Seq data from mouse and human cells, only genes with the same gene symbol in mouse and human transcriptomes were considered. After filtering out low-expression genes (mean expression (log2 RPKM) across the eight ESC to EpiLC time-points > 1.5, empirically derived from distribution of means), coefficient of variation for each gene was calculated as a measure of variability in gene expression, and the top 1000 genes with the highest variability in expression were used (Figures 7A and 7B).

### ChIP-Seq Data Analysis

Single-end 51 bp reads were mapped to the mouse (mm9) genome using Bowtie version 0.12.8 (Langmead et al., 2009), allowing up to two mismatches, retaining only reads that align to unique locations. For visualization on the UCSC Genome Browser and generation of screenshots, mapped reads were normalized to reads per million (RPM) and plotted as histograms using Bedtools version 2.26.0 (Quinlan, 2014). Enrichment of individual histone modifications (except for H3K9me2) or RNAPII at gene promoters (Figure 1C) was called based on normalized ChIP-Seq read density within the promoter region (±2 Kb of TSS) compared to input read density with the same region (>3-fold and FDR<0.01). For H3K9me2, given its broader footprint, ChIP-Seq read density within gene body instead of the promoter was used. For PCA unsupervised hierarchical clustering of histone modifications and RNAPII data (Figures S3A and S3F), ChIP-Seq signal for were quantile normalized to account for differences in signal-to-noise ratios.

### Correlation Analysis of Protein and mRNA expression

Global correlation between protein and mRNA expression for each time-point (Figure S6A) was calculated using Pearson correlation coefficient using only genes that were quantified at both the mRNA (log10 RPKM) and the protein level (log10 LFQ intensity). Global correlation between protein and mRNA fold-changes (compared to 0h data) for each time-point was calculated similarly (Figure S6B) and fitted using a local polynomial regression (Loess) model (Figure 4A). The correlation between mRNA and protein expression across the time-course, for individual genes, was calculated using Pearson correlation coefficient (Figures 4C, 4D, and S6D). To determine the relevance of each gene to ESCs and EpiLCs, we first rank-ordered the genes based on their mRNA (or protein) fold-changes in EpiLC (72h) vs ESC (0h) and then obtained the final rank-ordering of the genes (Figures 4D and S6C) based on the average of their two ranks (mRNA/protein expression-based).

### Comparative Analysis of Multi-ome Dynamics

Differentially regulated mRNAs, proteins and phosphosites at one or more time points (compared to 0h data) were determined using ANOVA test with multiple testing correction (FDR<0.01) (Figure S4). Volcano plots were used to visualize dynamic regulation by plotting the log2 fold change against –log10 of the permutation FDR adjusted *p*-value of the *t*-test on each mRNA, protein and phosphosite, respectively, at each time point. A scatter parameter of 0.1 (Tusher et al., 2001) which takes into account the log2 fold change and the statistical significance was used to identify dynamically regulated mRNAs, proteins, and phosphosites, respectively, at each time point. Percentage of dynamically regulated mRNAs, proteins, phosphosites and enriched H3K4me3 and H3K27me3 were calculated for each time point and scaled to the maximum percentage on transcriptome, proteome, phosphoproteome and epigenome layers. A local polynomial regression (Loess) was fitted to the scaled percentage calculated for each time point (Figure 2D). Magnitude of change for each mRNA, protein and phosphosite (Figure 2E) was determined by taking the highest absolute fold-change observed at all time points (compared to 0 h data): max(abs(*x*_*i*_/*x*_0_), *i* = 1 … *n*, where *x*_*i*_ (and *x*_0_) denotes the normalized value quantified at the *i*th (or 0 h) time-point for each mRNA, protein, or phosphosite.

### Gene Ontology Analysis

Gene Ontology (GO) analysis of differentially expressed genes (72h vs 0h; Figure 4B) was performed using only genes that were up- or down-regulated at both the mRNA and protein levels. To identify such genes, we integrated the proteomics and transcriptomics data using a previously published strategy (Yang et al., 2014) to group genes into eight classes based on the following criteria: (I) up-regulated at both the mRNA and protein levels, (II) up-regulated at the mRNA level but unchanged at the protein level, (III) up-regulated at the mRNA level but down-regulated at the protein level, (IV) unchanged at the mRNA level but down-regulated at the protein level, (V) down-regulated at both the mRNA and protein levels, (VI) down-regulated at the mRNA level but unchanged at the protein level, (VII) down-regulated at the mRNA level but up-regulated at the protein level, and (VII) unchanged at the mRNA level but up-regulated at the protein level. Class I (up-regulated) and class V (down-regulated) genes were analyzed for enriched GO categories (Figure 4B) using GO annotations (Gene Ontology, 2015).

### Kinase Activity Inference

To infer kinases active during ESC to EpiLC transition, we used CLUE (Yang et al., 2015), a fuzzy c-means clustering algorithm, to partition all phosphosites into 12 optimal clusters based on their temporal profiles (Figures S5A and S5B), and identified, for each cluster, kinases whose known substrates are enriched within that cluster. Known kinase-substrate relationships annotated in the PhosphoSitePlus database (Hornbeck et al., 2012) were used as a reference, and Fisher’s exact test was used to assess statistical significance of over-representation. Four out of the 12 clusters were found to be enriched for substrates with known kinases ERK/S6K/RSK, mTOR, p38a, and AKT (Figures 3A and 3B). An independent kinase perturbation analysis (Figure S5C) was performed using KinasePA (Yang et al., 2016b) to infer kinases active/regulated at various time-points during ESC to EpiLC induction, based on known kinase-substrate relationships annotated in PhosphoSitePlus database (Hornbeck et al., 2012).

### Pathway Enrichment Analysis

Pathway enrichment analysis (Figure 3C) was performed using the list of genes that encode for proteins containing the phosphosites from each of the inferred cluster. Pathway enrichment within a set of genes was evaluated by comparing that set of genes against genes within known pathways, as annotated in the Reactome database (https://reactome.org). Fisher’s exact test was used to assess statistical significance of over-representation.

### Substrate Prediction and Motif Analysis

Substrate prediction for ERK, mTOR, AKT, RSK/S6K and p38a was performed using an extended multiclass prediction version of the positive-unlabeled ensemble learning (Yang et al., 2016a). Briefly, the ensemble learning algorithm obtains the positive training instances by extracting known kinase-substrates from PhosphoSitePlus database and combines them with negative training instances obtained by randomly sampling from the rest of all identified phosphorylation sites using a positive-unlabeled learning technique. Throughout the training and prediction steps, the ensemble model integrates both the dynamic features extracted from time-resolved phosphoproteomics temporal profiles and the kinase recognition motif compiled from known substrates of each kinase and subsequently performs a multiclass classification to predict novel substrates for each kinase. Prediction results from the model were visualized as three-dimensional scatter plots with rainbow gradient colors from red to purple indicating most to least probable substrates of each kinase (Figure S5D, inset). Prediction results were also clustered to show their proximity to other predicted substrates of the same or a different kinase (Figure S5D). Consensus sequence motifs enriched within predicted substrates (Figure 3J) were identified using IceLogo (Colaert et al., 2009), using precompiled mouse Swiss-Prot sequence composition as the reference set.

## QUANTIFICATION AND STATISTICAL ANALYSIS

See Methods Details for details of quantification and statistical analysis.

## DATA AVAILABILITY

Mass spectrometry data generated for this study have been deposited to the ProteomeXchange Consortium (http://proteomecentral.proteomexchange.org/cgi/GetDataset), via the PRIDE (Deutsch et al., 2017) partner repository and will be available for access upon publication. RNA-Seq and ChIP-Seq data generated for this study have been deposited in the GEO repository under the accession number GSE117896 and will available for access upon publication.

